# A mechanism-annotated benchmark reveals limited fidelity to drug-response signatures in single-cell perturbation models

**DOI:** 10.64898/2026.08.19.745729

**Authors:** Lehang Li, Shaoming Duan, Xinyu Zha, Ye Fang, Yuhao Zhang, Xinyi Zhang, Yang Cao, Chuanyi Liu, Binxing Fang

## Abstract

Single-cell drug perturbation models are increasingly used to predict how compounds remodel cellular states, but they are still largely assessed by expression reconstruction. Whether high expression similarity reflects preservation of drug-response signatures remains unclear. Here we present scDrugPerturb-Bench, a mechanism-annotated benchmark that links matched control and drug-treated single-cell RNA-sequencing profiles to literature-curated directional key-gene evidence. The resource covers 181 datasets, 423 annotated response cases, 717 unique key genes and 2.5 million cells. We introduce the Mechanism Fidelity Score (MFS) to evaluate key-gene direction, effect-size recovery, gene-set coherence, mechanism specificity and pathway-level response polarity. Across 12 perturbation-prediction models, 3 baselines and 10 data splits, expression-similarity metrics were weakly aligned with MFS and selected different model configurations. Mechanism-aware selection improved early drug retrieval in a transcriptome-based drug design evaluation, indicating that MFS provides practical information beyond benchmark reporting. Systematic benchmarking revealed limited fidelity to drug-response signatures across cell-line and source-integrated settings. Frozen single-cell foundation model embeddings produced local, metric-dependent gains rather than universal improvements, and source context substantially reshaped model assessment. Hard-negative tests further showed that plausible perturbation responses can arise from non-specific transcriptional shortcuts. These results show that expression reconstruction is an insufficient proxy for preserving drug-response signatures and establish scDrugPerturb-Bench as a benchmark for mechanism-aware evaluation of single-cell drug perturbation models.

## 1. Introduction

Single-cell drug perturbation assays have made pharmacological responses measurable at cellular resolution. By pairing chemical perturbation with single-cell RNA sequencing, approaches such as sci-Plex [1], Perturb-seq-derived chemical screens and related assays map drug-induced transcriptional rewiring across cellular states, doses and contexts [2]. These data provide a foundation for computational perturbation models that could accelerate drug discovery. Such models could predict responses to candidate compounds, identify mechanism-specific biomarkers, nominate state-reversing drugs and prioritize therapeutic hypotheses before large-scale experimental screening. Yet experimental screens cannot exhaustively cover all combinations of drugs, doses, time points and cellular contexts. Whether these models preserve drug-response signatures, rather than fit perturbed expression distributions, remains unresolved.

A rapidly expanding set of perturbation-prediction models has been developed, including variational autoencoder-based models [3, 4], diffusion models [5, 6], transformer-based models [7, 8] and graph neural-network models [9]. In parallel, single-cell foundation models (scFM) [10–12] trained on large transcriptomic corpora are increasingly used as general-purpose representations for downstream prediction tasks. Existing benchmarks [13–18] have compared these methods across perturbation datasets and generalization settings, but they usually rely on expression-similarity metrics such as mean squared error, Pearson correlation, E-distance, maximum mean discrepancy and differential-gene overlap. These metrics are useful for assessing transcriptomic reconstruction, but high expression similarity does not imply that a model preserves the relevant drug-response signatures. In our analyses, a model could maintain high expression similarity in T0901317-treated human induced pluripotent stem cell-derived microglia while predicting the opposite direction for the LXR target genes *ABCA1, ABCG1* and *APOE* (Fig. 2a). This mismatch shows that current benchmarks leave a key evaluation gap: they do not directly test whether apparently accurate perturbation models preserve the relevant drug-response signatures. This gap is biologically consequential. Drug discovery rarely depends on global expression similarity alone [19, 20]. A useful model should identify which genes and pathways are altered by a compound, whether these changes occur in the correct direction, whether their effect sizes are biologically calibrated, whether functionally related genes respond coherently and whether the predicted response is specific to the drug-response signature rather than a generic stress, toxicity or cell-cycle programme [1, 21]. Models that miss these properties may nominate the wrong biomarkers, prioritize ineffective compounds, obscure target pathways or overstate confidence in virtual screening results [22]. Evaluating these properties requires case-specific annotations of the genes and pathways altered by each drug in each cellular context. Existing perturbation-prediction benchmarks or datasets rarely provide such mechanism annotations [13, 18].

Here we present scDrugPerturb-Bench, a mechanism-annotated benchmark designed to evaluate whether single-cell drug perturbation models preserve specific drug-response signatures. Starting from more than 50,000 PubMed records related to human drug perturbation and single-cell RNA-seq, we retained 93 studies and 181 datasets. The resulting resource spans five experimental sources, 137 cellular contexts, 101 drugs, 423 annotated response cases, 717 annotated unique key genes and 2.5 million cells. Each case links matched control and drug-treated single-cell RNA-seq profiles with curated drug, dose, time, cellular-context and directionally annotated key-gene evidence. This design converts drug-response statements dispersed across published studies into evaluation units that directly test whether predicted responses preserve the reported biology.

We introduce MFS, a mechanism-fidelity evaluation framework that measures complementary properties of predicted drug-response signatures. The Pattern Consistency Score evaluates whether key genes are predicted to change in the annotated direction. The Effect Size Recovery score measures whether predicted perturbation magnitudes match observed responses. The Gene-set Coherence Score assesses whether functionally related mechanism genes respond in a coordinated manner. The Mechanism Specificity Score tests whether predicted effects are enriched in mechanism-relevant genes rather than expression-matched non-specific genes. We further assess pathway-level response polarity using pathway Spearman correlation and pathway sign accuracy. By comparing MFS with conventional expression-similarity scores and transcriptome-based drug-retrieval [23] outcomes, we test whether mechanism-aware evaluation provides non-redundant information for assessing and selecting perturbation models.

Using scDrugPerturb-Bench, we evaluated 12 drug perturbation-prediction models and 3 baselines across 10 data splits spanning cell-line and source generalization. We further paired eight frozen scFMs and a PCA baseline with compatible downstream predictors to test whether pretrained representations improve mechanism fidelity. These analyses first show that expression similarity and mechanism fidelity are weakly coupled: models can generate plausible transcriptional responses while failing to preserve key-gene direction, pathway-level response or mechanism specificity. Transcriptome-based drug-retrieval experiments then show that mechanism-aware selection provides practical information for downstream representation choice, although it does not replace direct task-specific evaluation. Systematic benchmarking reveals limited fidelity to drug-response signatures in controlled cell-line settings and in heterogeneous source-integrated settings, where frozen scFM embeddings provide local, metric-dependent gains rather than universal improvements in mechanism fidelity. Source context further reshapes mechanism-fidelity profiles, model rankings and the identity of preferred pipelines. Finally, hard-negative analyses show that apparent signature preservation can arise from non-specific transcriptional shortcuts, including generic-response, signature-specificity and global-strength effects. Together, these findings show that single-cell drug perturbation models should be evaluated not only by how closely they reconstruct expression profiles, but also by whether they preserve the case-specific signatures that make predictions biologically actionable.

## 2. Results

### 2.1. A mechanism-annotated benchmark for single-cell drug perturbation prediction

#### Datasets

Existing single-cell drug perturbation benchmarks rarely annotate the experimentally supported mechanism of each drug response. Such information is usually dispersed across article text, figures and supplementary materials, rather than stored with the expression matrix. We therefore built a literature-to-matrix curation workflow that combines PubMed retrieval, key-insight extraction, automated single-cell RNA-seq processing, manual verification and data cleaning (Fig. 1a). This workflow extracted the cellular context, drug identity, dose, key post-perturbation gene changes and the source evidence supporting each annotation. Candidate cases were first assembled through a semi-automated curation process and then manually verified against the original publication. We further checked whether the measured expression matrix reproduced the reported direction of the key gene changes; cases with inconsistent literature evidence and measured expression responses were excluded. All retained data were processed through a unified workflow for cell filtering, gene mapping, gene filtering and normalization. The final benchmark includes matched control matrices, drug-perturbed matrices, drug metadata and curated annotations of mechanism-relevant gene changes.

**Figure 1.**
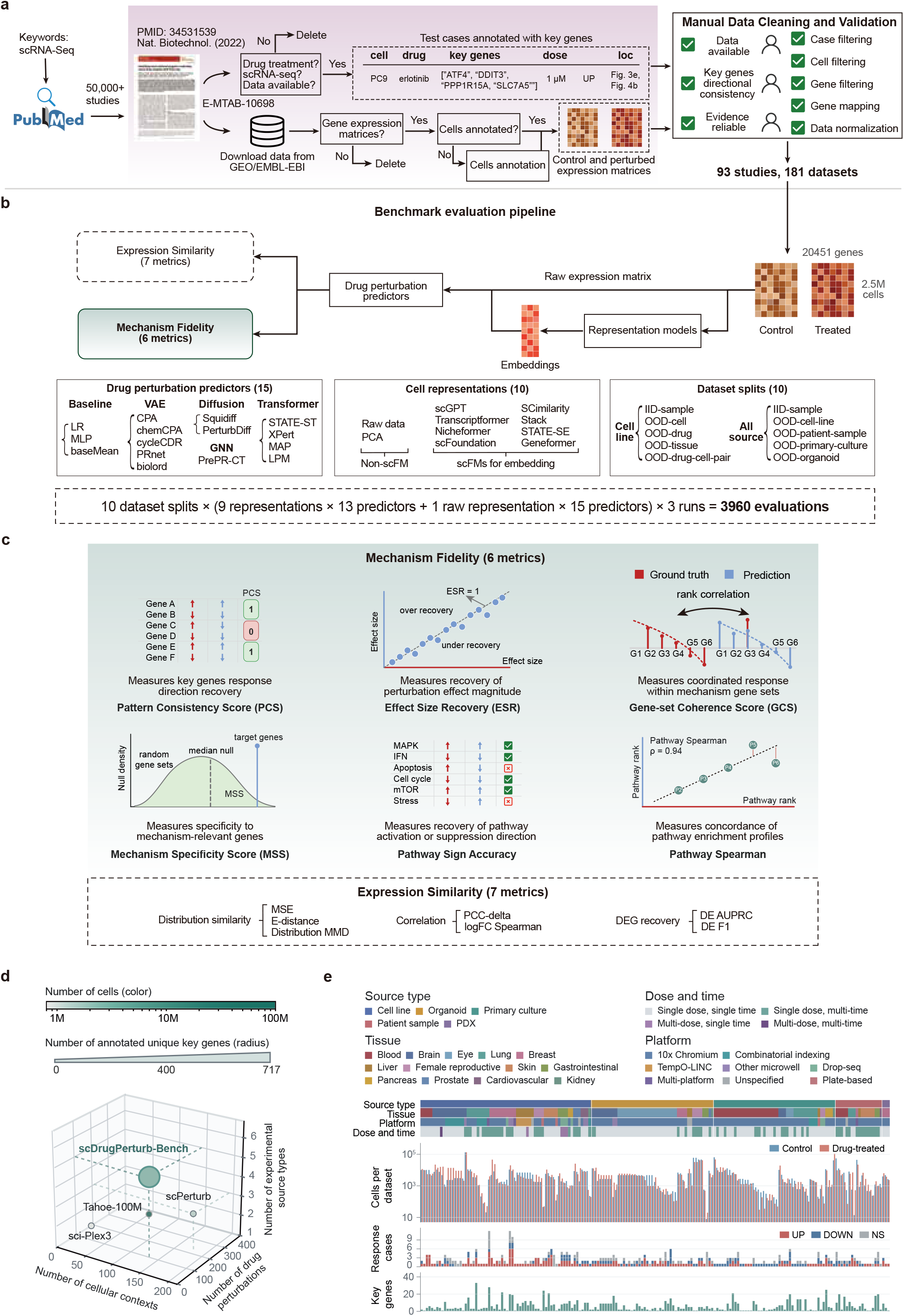
Literature-to-benchmark workflow and mechanism-aware evaluation of single-cell drug perturbation models. **a**, Data-collection workflow. Candidate studies were retrieved from PubMed using keyword-based search, screened for drug perturbation and single-cell RNA-seq relevance, processed through key-insight extraction and automated single-cell data processing, and finalized by manual verification, data cleaning and quality control. **b**, scDrugPerturb-Bench evaluation pipeline. Across ten data splits, processed datasets were represented as raw expression matrices or as embeddings from nine representation methods, and then evaluated with 12 perturbation-prediction models and three baselines. **c**, Mechanism-fidelity and expression-similarity metrics used in scDrugPerturb-Bench. **d**, Benchmark comparison with representative perturbation resources across cellular contexts, perturbation coverage and experimental source types. **e**, Dataset-level composition of scDrugPerturb-Bench across experimental sources, tissue groups, profiling platforms, dose–time designs, response calls and annotated key genes.

**Figure 2.**
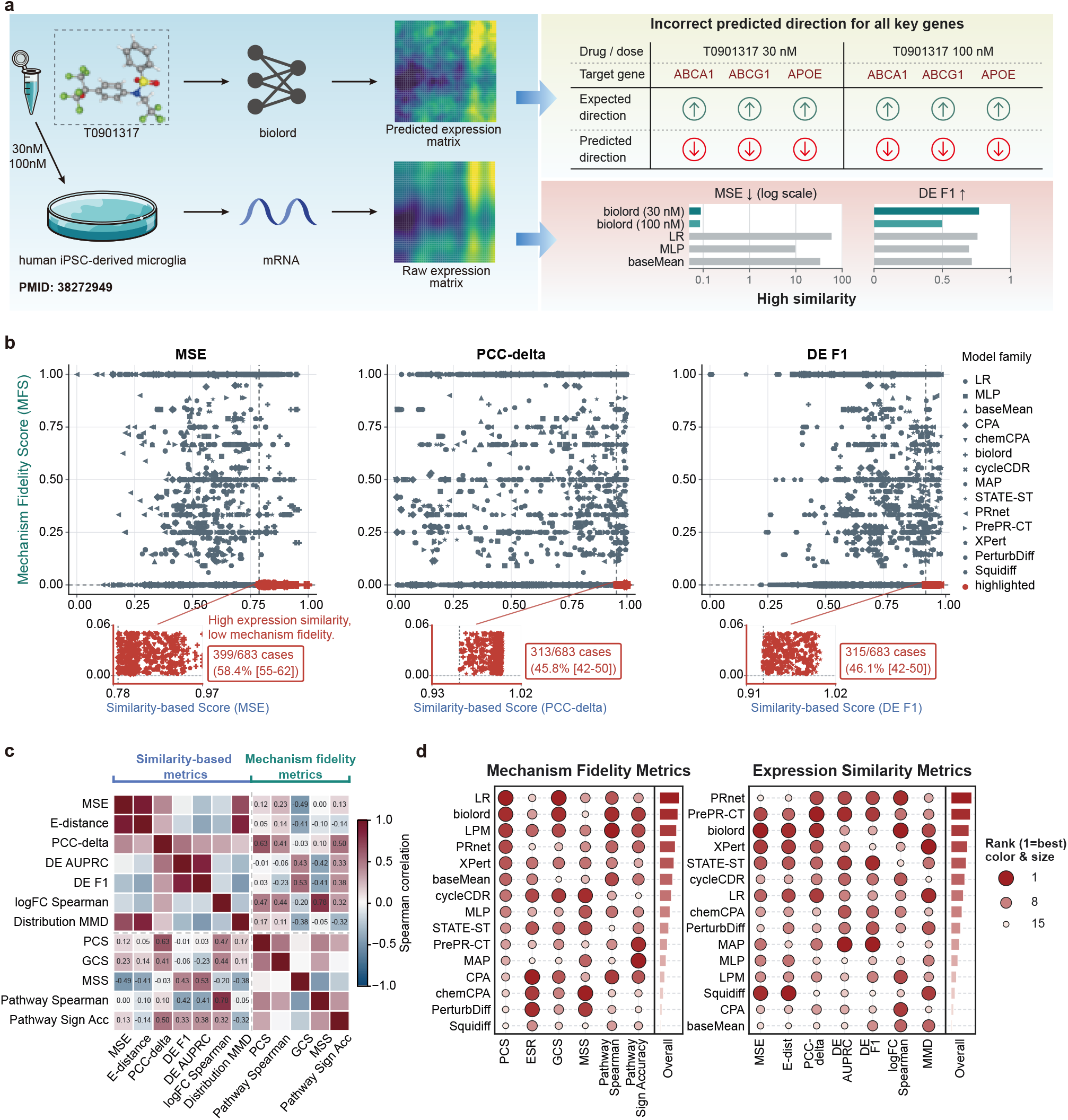
Expression similarity is insufficient for assessing mechanism fidelity. **a**, Mechanistic case study in T0901317-treated human iPSC-derived microglia, where high expression similarity coincided with reversed prediction of 3 upregulated LXR target genes. **b**, Discordant high-expression-similarity and low-mechanism-fidelity cases, defined by top-20% direction-oriented similarity-based score and bottom-20% PCS in the iid-sample setting. **c**, Spearman correlations between expression similarity metrics and mechanism fidelity metrics in the iid-sample setting after orienting all metrics. **d**, Discordance between model rankings induced by ESS and PCS-prioritized MFS across cell-line data-splitting settings.

Starting from more than 50,000 PubMed records related to human drug perturbation and single-cell RNA-seq, this workflow retained 93 studies and 181 datasets comprising 2.5M cells. The resulting resource spans five experimental sources, 137 cellular contexts, 101 drugs, 13 tissue groups and 717 annotated unique key genes (Fig. 1d,e and Supplementary Fig. S1). At the dataset level, the benchmark includes cell lines (66 datasets), primary cultures (47), organoids (47), patient samples (18) and patient-derived xenografts (3). Most datasets were generated with 10x Chromium (135 datasets), with additional coverage from combinatorial-indexing, Drop-seq, TempO-LINC, microwell, plate-based and multi-platform assays. Experimental designs ranged from single-dose, single-time measurements to multidose and multitime studies. Across these datasets, we curated 423 response cases, including 195 upregulated, 126 downregulated and 102 non-significant responses. This coverage was intended to preserve the biological and technical heterogeneity encountered in real perturbation studies while ensuring that each benchmark case has explicit evidence for mechanism-aware evaluation. We designed scDrugPerturb-Bench as a versioned resource that can be maintained and updated as new single-cell drug perturbation datasets and mechanism annotations become available.

Compared with existing single-cell drug perturbation resources, the main advantage of scDrugPerturb-Bench is not maximal cell scale alone, but the coupling of multi-source and multi-context perturbation profiles with manually curated mechanism-level information (Fig. 1d,e). sci-Plex3[1], scPerturb[18] and Tahoe-100M[24] provide important expression-profile resources for perturbation modelling, but they lack case-specific directional response annotations for mechanism-aware model evaluation. scDrugPerturb-Bench links heterogeneous perturbation matrices to curated drug, dose, time, cellular-context and key-gene evidence, allowing model assessment to move beyond transcriptional prediction accuracy toward mechanism consistency, key-gene directionality and interpretation of perturbation processes.

#### Dataset splits

We evaluated models under ten data splits that reflect major sources of heterogeneity in single-cell drug-response prediction (Fig. 1b). These splits spanned two complementary generalization regimes. In the cell-line generalization regime, we considered iid-sample, OOD-drug, OOD-cell, OOD-tissue and OOD-drug-cell-pair settings. The iid-sample split divides cells within each dataset into training and test sets, so that the same drug and cellular context are represented in both. By contrast, OOD-drug, OOD-cell and OOD-tissue splits hold out drugs, cell types or tissues from training, and the OOD-drug-cell-pair split holds out specific drug–cell combinations. In the source generalization regime, we evaluated whether models trained on one set of experimental sources could predict responses in previously unseen source domains. Patient-derived xenograft (PDX) data were retained only as a training source because this category contained too few datasets for a standalone test split.

#### Methods

We evaluated two perturbation-prediction pipelines (Fig. 1b). In the direct pipeline, processed expression matrices were provided to each perturbation-prediction model and model outputs were compared on matched held-out perturbation cases. In the representation-augmented pipeline, the same expression matrices were first converted into cell embeddings by an upstream representation model and these embeddings were then used as inputs to compatible downstream perturbation-prediction models. Across these pipelines, the benchmark comprised ten input data types: raw expression matrices and embeddings from nine representation methods. This design allowed us to test both the standalone performance of perturbation predictors and whether fixed transcriptomic representations improve drug-response prediction under the same data splits and evaluation metrics.

The perturbation-prediction benchmark comprised 12 published models and three baseline predictors. The baselines were linear regression[25], MLP[25] and a mean-response predictor[25]. The published models spanned VAE-based approaches, including CPA[3], chemCPA[4], cycleCDR[26], PRnet[27] and biolord[28]; transformer-based approaches, including STATE-ST[29], XPert[7], MAP[30] and LPM[8]; diffusion-based approaches, including Squidiff[5] and PerturbDiff[6]; and the graph neural network-based model PrePR-CT[9]. For the representation-augmented pipeline, we evaluated eight single-cell foundation models (scFMs): scGPT[31], scFoundation[11], Geneformer[12], Transcriptformer[32], Nicheformer[33], SCimilarity[34], Stack[35] and STATE-SE[29], with PCA included as a non-neural representation baseline. scFMs were used only as frozen feature extractors and were not fine-tuned on the benchmark. Thus, differences in downstream performance reflect the utility of the extracted cell representations rather than task-specific adaptation of the foundation models.

#### Evaluation

Together, the framework produced 4,500 scored evaluations across 12 perturbation-prediction models, three baselines, ten data splits and ten cell representations (Fig. 1b). We evaluated predictions with seven conventional expression-similarity metrics and six MFS metrics (Fig. 1c). MFS integrates six complementary metrics that decompose mechanism fidelity into gene-level and pathway-level properties. At the gene level, Pattern Consistency Score (PCS) measures directional consistency of curated response genes, Effect Size Recovery (ESR) measures agreement in response magnitude, Gene-set Coherence Score (GCS) measures preservation of coordinated responses within annotated gene sets and Mechanism Specificity Score (MSS) measures whether predicted effects are enriched in mechanism-relevant genes rather than diffuse background changes. At the pathway level, Pathway Spearman measures concordance between predicted and observed enrichment profiles, whereas Pathway Sign Accuracy measures whether significantly perturbed pathways are assigned the correct activation or suppression direction. Whenever MFS was used for ranking or selection, we used a PCS-prioritized rule that treated PCS as the primary ordering criterion and used gene-set and pathway-level metrics to resolve near ties (Methods).

In parallel, the expression similarity score (ESS) metrics in Fig. 1c quantified how closely predicted perturbed profiles matched measured perturbed profiles. These included distributional metrics, MSE, E-distance and Distribution MMD; correlation-based metrics, PCC-delta and logFC Spearman; and differential-expression agreement metrics, DE AUPRC and DE F1. This paired evaluation design allowed us to test whether expression-level similarity is sufficient for selecting models with high mechanism fidelity, and whether MFS provides non-redundant information for model assessment and downstream model selection.

### 2.2. Expression similarity does not imply mechanism fidelity

Drug perturbation prediction models are commonly evaluated by similarity between predicted and measured expression profiles. This criterion is necessary for assessing transcriptomic reconstruction, but it does not test whether a model preserves the drug-response signature that makes a prediction biologically actionable. A representative case illustrates this gap (Fig. 2a). In human induced pluripotent stem cell (iPSC)-derived microglia treated with T0901317, the measured response showed upregulation of the LXR target genes *ABCA1, ABCG1* and *APOE*. biolord nevertheless predicted the opposite direction for all 3 genes at both 30 nM and 100 nM, yielding PCS values of 0 while maintaining high expression-similarity scores. Thus, an apparently accurate expression profile failed the central signature-level criterion: preservation of the direction of a known drug-response signature.

We therefore compared conventional expression-similarity metrics with mechanism-fidelity metrics that evaluate key-gene direction, gene-set structure, mechanism specificity and pathway-level response polarity. This analysis was performed in the cell-line generalization regime across five data splits: iid-sample, OOD-drug, OOD-cell, OOD-tissue and OOD-drug-cell-pair. High expression similarity frequently concealed limited fidelity to drug-response signatures at the case level (Fig. 2b and Supplementary S3). For these case-level discordance analyses, error and distance metrics were displayed as direction-oriented similarity-based scores rather than as raw values. MSE and E-distance were log-transformed and reversed after min–max normalization, whereas Distribution MMD was reversed after min–max normalization without log transformation. Thus, larger values in Fig. 2b consistently indicate higher expression similarity, including lower raw MSE, E-distance or Distribution MMD. We defined discordant cases as predictions in the top 20% for the direction-oriented expression-similarity score but the bottom 20% for Pattern Consistency Score. Such cases occurred in every split. In iid-sample, 34.3–59.4% of high-expression cases also had low PCS, including 58.4% for MSE and 59.4% for E-distance. Discordance was strongest under OOD-cell generalization: MSE, E-distance, Distribution MMD and PCC-delta showed high-expression/low-PCS rates of 99.1%, 91.8%, 97.3% and 89.1%, respectively. OOD-tissue and OOD-drug-cell-pair also showed substantial discordance, with ranges of 47.7–66.4% and 45.3–76.9%, respectively, whereas OOD-drug still reached 20.0–49.1%. This pattern shows that a model can place predicted cells close to the measured perturbed distribution while still failing to preserve the direction of curated response genes.

Expression similarity and mechanism fidelity also provided largely non-redundant information at the metric level (Fig. 2c and Supplementary Fig. S2a). After orienting all metrics so that higher values indicated better performance, expression–mechanism correlations were weak across the five data-splitting settings: across 175 cross-metric comparisons, the median Spearman correlation was 0.052 and the median absolute correlation was 0.241. In total, 58.9% of absolute correlations were below 0.3 and 88.6% were below 0.5. This weak coupling was most pronounced in OOD-cell, where the median absolute expression–mechanism correlation was 0.136, and remained limited even in OOD-drug, the most aligned split, where the median absolute correlation was 0.335. Individual metric pairs could be strongly associated, but the direction was not stable across settings. For example, logFC Spearman and Pathway Spearman were positively correlated in iid-sample (*ρ* = 0.777), whereas DE AUPRC and Pathway Spearman were strongly anticorrelated in OOD-drug (*ρ* = − 0.766). Thus, selected differential-expression metrics captured part of the responsive-gene signal, but they did not establish whether the specific literature-curated genes and pathways were preserved with the correct direction and specificity.

The weak metric coupling translated into different model rankings (Fig. 2d and Supplementary Fig. S2b). The PCS-prioritized MFS and ESS never selected the same top-ranked model across the five splits, and their overall rank correlation was weak (median Spearman *ρ* = −0.104; mean *ρ* = −0.030). In iid-sample, MFS ranked linear regression first whereas ESS ranked PRnet first. In OOD-drug, OOD-tissue and OOD-drug-cell-pair, MFS ranked LPM first, whereas ESS ranked MAP, STATE-ST and STATE-ST first, respectively. These rank reversals show that model choice depends strongly on whether the evaluation target is expression reconstruction, key-gene direction preservation, mechanism specificity or pathway-level response.

Together, these analyses show that expression similarity is an insufficient proxy for preserving drug-response signatures. Mechanism fidelity metrics provide non-redundant information about response direction, pathway structure and mechanism specificity, and are therefore necessary for assessing whether virtual cell models capture biologically meaningful drug responses.

### 2.3. Mechanism-aware representation selection improves early drug retrieval

This separation between expression similarity and mechanism fidelity motivated an application-level test of whether a PCS-prioritized MFS ranking improves representation selection beyond benchmark reporting. Downstream transcriptome-based drug retrieval was used as an external validation setting rather than as an independent retrieval benchmark. The rationale was that a representation supporting mechanism-faithful perturbation prediction should allow the true perturbing drug to be retrieved earlier and more accurately. We evaluated eight frozen single-cell foundation models (scFMs) and PCA across four out-of-distribution settings in the cell-line generalization scenario (Fig. 3a). The workflow linked two tasks. First, chemCPA[4] predicted held-out drug responses from each fixed representation, and these predictions were scored and ranked by PCS-prioritized MFS Overall and ESS Overall. Second, the predicted transcriptomic response was supplied to CURE[23], which generated a query drug fingerprint and ranked candidate compounds. Drug-retrieval performance was summarized as DSS Overall, which combined early Hit@K, early NDCG@K and mean reciprocal rank. This setup provided a matched test of whether representations selected by PCS-prioritized MFS or ESS also improved retrieval of the true perturbing drug.

**Figure 3.**
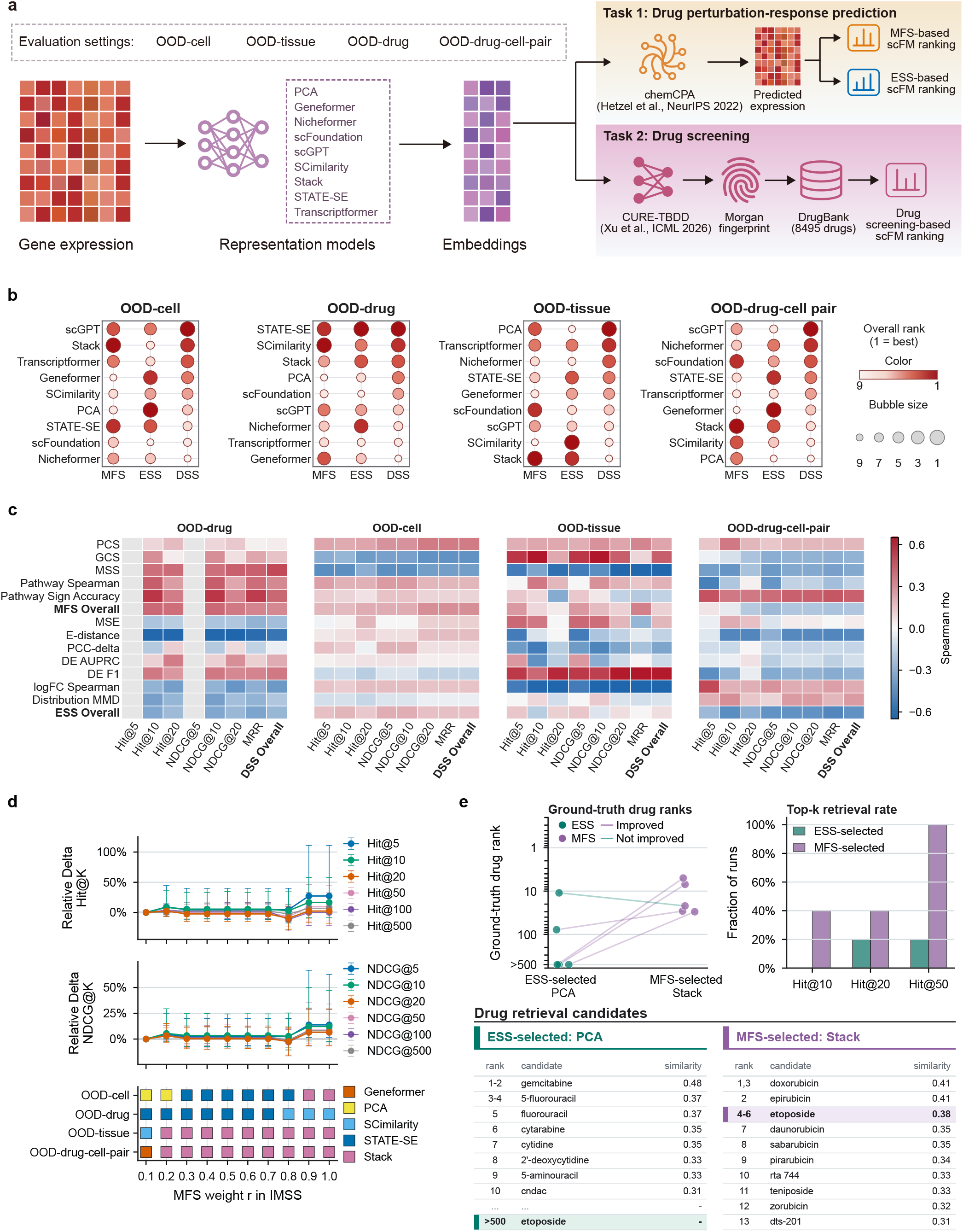
Mechanism-guided representation selection prioritizes effective drugs in downstream retrieval. **a**, Experimental workflow comparing PCS-prioritized MFS- and ESS-based representation selection across embedding-informed perturbation prediction and CURE-based drug retrieval. **b**, Representation-ranking comparison across mechanism fidelity, expression similarity and downstream drug-retrieval performance in the OOD setting. **c**, Spearman correlations between perturbation-prediction metrics and downstream drug-retrieval scores in the OOD setting. **d**, Sensitivity analysis of the integrated representation-selection score, which combines PCS-prioritized MFS and ESS rank percentiles. **e**, OOD-cell etoposide case comparing ESS-selected PCA and PCS-prioritized MFS-selected Stack in CURE-based drug retrieval.

PCS-prioritized MFS, ESS and downstream retrieval rankings were not interchangeable (Fig. 3b). Agreement was highest in OOD-drug: STATE-SE ranked first by both ESS and DSS and second by

MFS, whereas SCimilarity ranked first by MFS and second by DSS. In OOD-cell, the MFS-selected representation Stack ranked second by DSS, whereas the ESS-selected representation PCA ranked only sixth by DSS. Conversely, scGPT ranked first by DSS but third by MFS and fourth by ESS. The disconnect was stronger in OOD-tissue and OOD-drug-cell-pair. In OOD-tissue, PCA ranked first by DSS but ninth by ESS, whereas Stack ranked first by MFS and second by ESS but ninth by DSS. In OOD-drug-cell-pair, scGPT ranked first by DSS despite ranking fifth by MFS and eighth by ESS. Across all representations and splits, Spearman correlation with DSS was 0.096 for MFS and -0.008 for ESS.

Correlation analysis gave a similar picture (Fig. 3c). MFS Overall was more consistently associated with DSS Overall than ESS Overall, but the correlations were modest and context dependent. PCS correlated positively with DSS in all four settings, with a mean Spearman correlation of 0.203 and a maximum of 0.356 in OOD-cell. Pathway Sign Accuracy was positive in three settings, with a mean correlation of 0.196 and a maximum of 0.433 in OOD-drug-cell-pair. MFS Overall was positive in OOD-cell, OOD-drug and OOD-tissue (*ρ* = 0.300, 0.317 and 0.110), but negative in OOD-drug-cell-pair (*ρ* = −0.217). ESS Overall averaged -0.127 and was negative in OOD-drug and OOD-drug-cell-pair (*ρ* = −0.289 and -0.451). Individual expression metrics were less stable: DE F1 showed the strongest positive association with DSS in OOD-tissue (*ρ* = 0.642), whereas logFC Spearman was strongly negative in the same split (*ρ* = −0.822). These descriptive correlations showed that mechanism-aware metrics provided complementary selection information, but did not replace direct DSS evaluation.

Combining PCS-prioritized MFS and ESS improved representation selection for drug retrieval when MFS was weighted more heavily (Fig. 3d). Increasing the MFS weight changed the selected representation in several splits and produced the largest early-retrieval gains when *r* = 0.9 or *r* = 1.0. At these weights, the IMSS-selected representations were identical to the MFS-selected representations and improved Hit@5, Hit@10, Hit@50 and Hit@500 by 27.3%, 16.7%, 9.3% and 7.0%, respectively, relative to ESS-selected representations. NDCG showed similar gains, with increases of 13.8%, 12.2%, 8.8% and 8.1% for NDCG@5, NDCG@10, NDCG@50 and NDCG@500, respectively. The clearest bootstrap-supported effects were concentrated at Top-5, where the 95% confidence intervals for Hit@5 and NDCG@5 did not cross zero. The trend was not monotonic, however: *r* = 0.8 triggered a representation switch in OOD-drug and reduced several Hit@K and NDCG@K values.

The same pattern was visible in a representative OOD-cell case (Fig. 3e). For etoposide, ESS selected PCA, whereas PCS-prioritized MFS selected Stack. PCA retrieved etoposide poorly across five runs, with ranks *>* 500, *>* 500, 78, *>* 500 and 11. Stack ranked etoposide within the top 50 in all five runs, with ranks 7, 30, 29, 5 and 22. This changed Hit@10 from 0% to 40%, Hit@20 from 20% to 40% and Hit@50 from 20% to 100%. In the representative run, PCA prioritized several antimetabolite or nucleoside-like compounds, whereas Stack ranked etoposide 4-6th and placed related topoisomerase-II-associated drugs, including doxorubicin, epirubicin, daunorubicin and teniposide, near the top of the candidate list. This case illustrates how mechanism-aware selection can improve early retrieval while producing a more mechanism-relevant candidate neighbourhood.

### 2.4. Cell-line benchmarks reveal limited fidelity to drug-response signatures

We next asked whether current predictors and frozen single-cell foundation model (scFM) embeddings preserved annotated drug-response signatures in a controlled cell-line regime (Fig. 4a). Across five cell-line splits, we compared Raw input, PCA and eight frozen scFM representations coupled to 12 perturbation-prediction models and three baselines. We then assessed absolute mechanism fidelity, ranking stability, representation-specific gains, selection robustness and computational cost.

**Figure 4.**
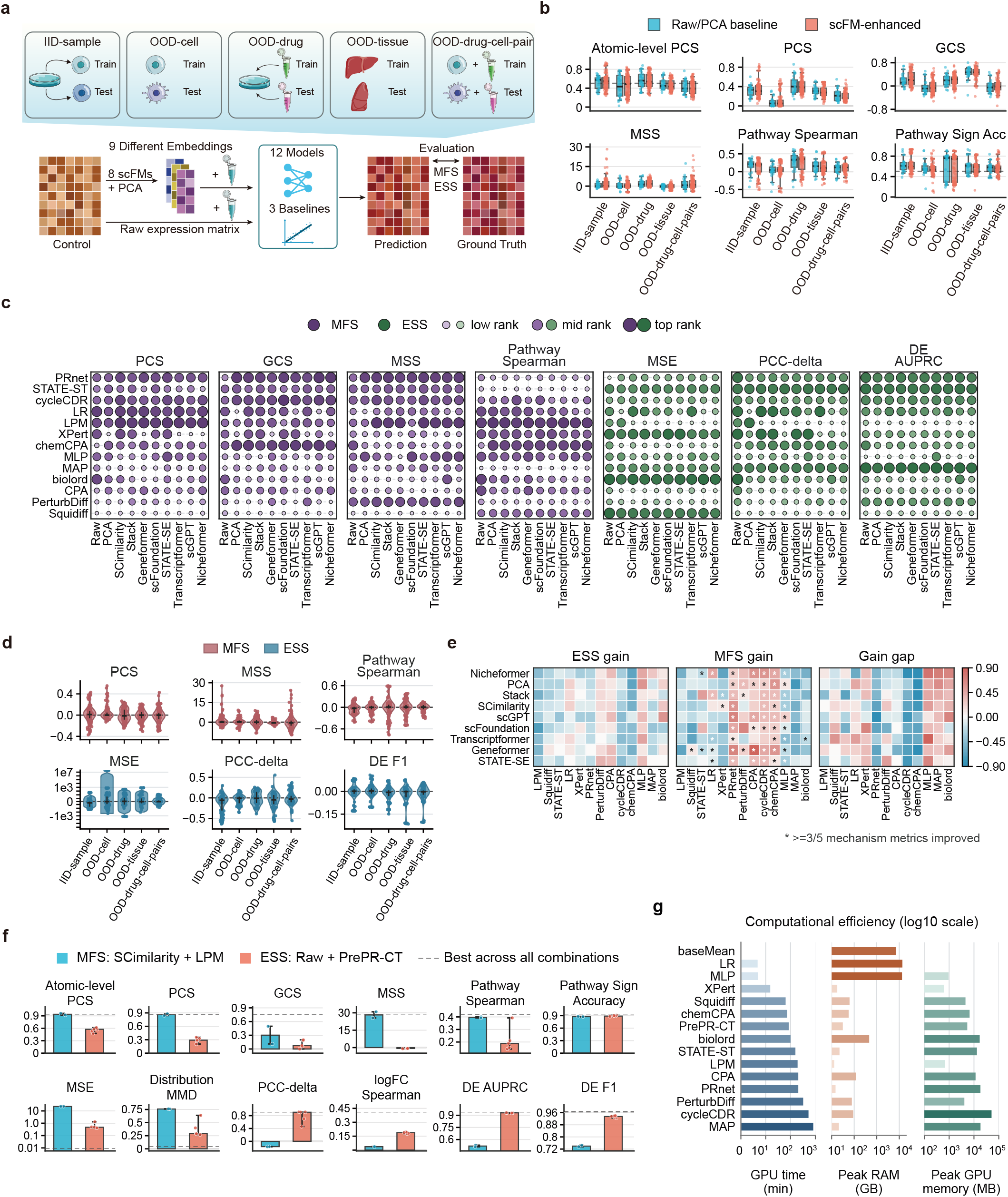
Cell-line perturbation benchmarks reveal limited fidelity to drug-response signatures, context-dependent representation gains and computational trade-offs. **a**, Cell-line benchmarking design across iid-sample, OOD-cell, OOD-drug, OOD-tissue and OOD-drug-cell-pair splits. Raw expression matrices, PCA embeddings and frozen scFM embeddings were coupled to perturbation-prediction models and evaluated by MFS and ESS. **b**, Pipeline-level distributions of six mechanism-fidelity components across the five cell-line splits, comparing Raw/PCA baseline pipelines with scFM-enhanced pipelines. **c**, IID-sample ranking landscape for predictor–representation combinations across representative MFS and ESS metrics. Dot size and colour intensity indicate relative rank within each metric. **d**, Direction-oriented gains of PCA and scFM embeddings relative to matched Raw-input predictors across five cell-line splits, shown for representative mechanism-fidelity and expression-similarity metrics. **e**, IID-sample gain maps for each embedding–predictor combination, summarizing ESS gain, PCS-prioritized MFS gain and the gain gap, defined as ESS gain minus MFS gain, relative to matched Raw-input predictors. Asterisks mark combinations improving at least three of five mechanism metrics. **f**, Metric-wise performance of the best iid-sample combinations selected by PCS-prioritized MFS or ESS. Dashed lines mark the best score achieved by any combination for each metric. **g**, Practical resource requirements of Raw-input perturbation-prediction models in iid-sample, measured by GPU runtime, peak RAM and peak GPU memory on log-scaled axes.

Absolute mechanism fidelity was limited across cell-line settings (Fig. 4b). Atomic-level PCS and Pathway Sign Accuracy were often above their reference levels, indicating partial preservation of gene-direction and pathway-polarity signals. However, GCS, MSS and Pathway Spearman remained low in several OOD settings. These metrics showed weaker preservation of coordinated key-gene structure, mechanism specificity and pathway-response ordering. Frozen scFM embeddings did not shift all mechanism dimensions in the same direction. MSS improved most consistently across splits, including from approximately -0.33 to 0.26 in OOD-cell and from 1.34 to 1.86 in OOD-drug. GCS increased or remained similar, including a shift from 0.12 to 0.21 in iid-sample and from 0.19 to 0.23 in OOD-drug. By contrast, Pathway Spearman improved in OOD-cell, OOD-drug and OOD-drug-cell-pair, but decreased in iid-sample and OOD-tissue. Atomic-level PCS improved most clearly in OOD-cell, whereas PCS and Pathway Sign Accuracy did not show a systematic scFM advantage. Thus, frozen scFM embeddings redistributed mechanism-fidelity performance across response dimensions rather than producing a uniform gain.

The highest-ranked pipelines changed across both metrics and data splits (Fig. 4c and Supplementary Fig. S5a). In iid-sample, high-ranking mechanism-fidelity combinations included SCimilarity–LPM, PCA–LPM, scFoundation–cycleCDR and scGPT–MAP. Raw-input combinations remained competitive for expression-similarity metrics, including Raw–biolord for MSE, PCC-delta and logFC Spearman, and Raw–MAP for DE AUPRC and DE F1. In OOD-cell, the leading MFS combinations shifted toward chemCPA paired with Geneformer, Nicheformer, scGPT or SCimilarity. In OOD-drug, OOD-tissue and OOD-drug-cell-pair, MLP paired with selected scFM or PCA representations became prominent among high-ranking MFS pipelines.

No representation was consistently dominant. Stack, SCimilarity and scFoundation ranked highly in iid-sample, Geneformer was most prominent in OOD-cell, and SCimilarity and scGPT performed well in OOD-drug. scGPT was relatively strong in OOD-tissue and OOD-drug-cell-pair. Raw and PCA pipelines also remained among the leading mechanism-fidelity results in parts of OOD-drug and OOD-tissue. Across pipeline-level rankings, the Spearman correlations between MFS and ESS were low in most splits: 0.03 in iid-sample, 0.32 in OOD-cell, -0.13 in OOD-drug, 0.02 in OOD-tissue and 0.09 in OOD-drug-cell-pair. These rank reversals show that expression reconstruction, key-gene directionality and pathway-level mechanism fidelity selected different model configurations.

Direct comparison with Raw input confirmed that frozen scFM embeddings did not yield broad, metric-consistent gains (Fig. 4d and Supplementary Fig. S4). Across paired comparisons, scFM effects were distributed around zero and frequently changed sign across metrics. In iid-sample, PCS showed a small positive shift, with about 60% of scFM–predictor combinations exceeding their matched Raw-input pipeline. However, MSS was close to unchanged and Pathway Spearman decreased. In OOD-cell, MSS showed the clearest improvement, with a median gain of about 0.22, whereas PCS and Pathway Spearman remained close to zero. In OOD-drug, Pathway Spearman increased slightly, but PCS decreased and MSS showed little overall change. In OOD-tissue and OOD-drug-cell-pair, Pathway Spearman showed modest positive shifts, but MSS declined, with median gains of approximately -0.54 and -0.40, respectively. Expression-similarity metrics showed the same lack of coordination. MSE improved slightly in OOD-cell and OOD-drug, but PCC-delta and DE F1 did not improve consistently. These paired analyses argue against treating frozen scFM embeddings as a universal replacement for Raw expression input.

Representation gains were instead highly specific to the embedding and predictor used together (Fig. 4e and Supplementary Fig. S5b). The gain gap was defined as ESS gain minus MFS gain. Positive values therefore indicate larger expression-similarity gains, whereas negative values indicate larger mechanism-fidelity gains. Across 117 representation–predictor combinations summarized over five splits, the median ESS gain and median PCS-prioritized MFS gain were both approximately -0.11. Thus, adding a representation did not usually outperform the matched Raw-input pipeline. The exceptions were strongly predictor dependent. PRnet, cycleCDR and chemCPA achieved median MFS gains of about 0.33 when paired with selected embeddings, whereas CPA and PerturbDiff had median MFS gains of about 0.22. In contrast, MLP, XPert and MAP showed negative MFS gains overall, even when expression gain was positive for some MLP combinations. Individual combinations also separated mechanism and expression benefits. Geneformer–CPA showed concurrent gains in MFS (0.89) and ESS (0.33), and Nicheformer–CPA, scGPT–CPA and scFoundation–PerturbDiff also improved both objectives. By contrast, Geneformer–chemCPA, Nicheformer–chemCPA and scGPT– PRnet increased MFS despite negative ESS gain, whereas Transcriptformer–MLP, Nicheformer–MLP and scGPT–biolord increased ESS while decreasing MFS. The overall Spearman correlation between ESS gain and MFS gain was approximately -0.04, indicating that representation-induced expression improvement did not predict improved drug-response signature fidelity.

We next evaluated whether the best combinations selected by PCS-prioritized MFS or ESS remained optimal across other mechanism-fidelity and expression-similarity metrics (Fig. 4f and Supplementary Fig. S6). In iid-sample, MFS selected SCimilarity–LPM, whereas ESS selected Raw–PrePR-CT. The MFS-selected pipeline reached or approached the best observed values for Atomic-level PCS, PCS, GCS, MSS and Pathway Spearman, including Atomic-level PCS of 0.940, PCS of 0.860 and MSS of 28.225. The ESS-selected pipeline was stronger across expression metrics, including MSE of 0.476, PCC-delta of 0.908, DE AUPRC of 0.931 and DE F1 of 0.935. Their global ranks were nearly reversed: SCimilarity–LPM ranked first by MFS but 125th by ESS, whereas Raw–PrePR-CT ranked first by ESS but 76th by MFS.

The OOD analyses showed the same separation in selection objective. In OOD-cell, MFS selected Geneformer–MAP, whereas ESS selected Raw–PrePR-CT. In OOD-drug and OOD-tissue, MFS selected Raw–LPM, whereas ESS selected Raw–MAP and Raw–STATE-ST, respectively. In OOD-drug-cell-pair, MFS selected Raw–LPM and ESS selected STATE-SE–STATE-ST. Across these splits, MFS-selected pipelines generally improved key-gene direction or mechanism-specific metrics, but they did not dominate all mechanism components. Conversely, ESS-selected pipelines retained stronger expression reconstruction while showing weaker Atomic-level PCS or PCS. These comparisons show that mechanism-aware and expression-aware criteria optimize different objectives, and that cross-split performance still requires explicit OOD evaluation rather than inference from a single split.

Resource profiling showed that benchmark practicality must be considered together with predictive performance (Fig. 4g). In the Raw-input iid-sample setting, lightweight baselines and compact predictors, including the mean predictor, linear regression, chemCPA, MLP and XPert, had comparatively low GPU runtimes. Larger generative or sequence models required longer runs, with MAP showing the largest GPU-time burden. The dominant resource bottleneck differed by method: biolord and CPA were constrained mainly by peak RAM, whereas MAP, cycleCDR, biolord, PRnet and STATE-ST required higher peak GPU memory. Thus, scalable use of a benchmark requires reporting runtime, RAM and GPU memory alongside MFS and ESS, because a strong score on one metric may not translate into a practical high-throughput workflow.

Together, the cell-line analysis shows that mechanism fidelity and representation choice must be evaluated jointly. scFM embeddings did not provide a universal gain in mechanism fidelity, mechanism-aware selection did not improve all mechanism-fidelity dimensions, and some complex predictors imposed substantial runtime or memory costs. These results establish the baseline limitation in a relatively controlled cell-line regime.

### 2.5. Source context reshapes mechanism fidelity in multi-source perturbation benchmarks

Cell-line benchmarks provide a controlled regime for evaluating drug-, cell- and tissue-level generalization. However, single-cell drug perturbation data are increasingly assembled across heterogeneous experimental systems, including cell lines, primary cultures, organoids and patient samples. We therefore analysed a distinct source-integrated regime to test whether mechanism fidelity, representation gains and model rankings remained stable when source context, input representation and predictor choice varied jointly (Fig. 5a). Across five source-aware splits, we compared Raw expression input, PCA and eight frozen scFM representations coupled to perturbation-prediction models and baselines using both MFS and ESS.

**Figure 5.**
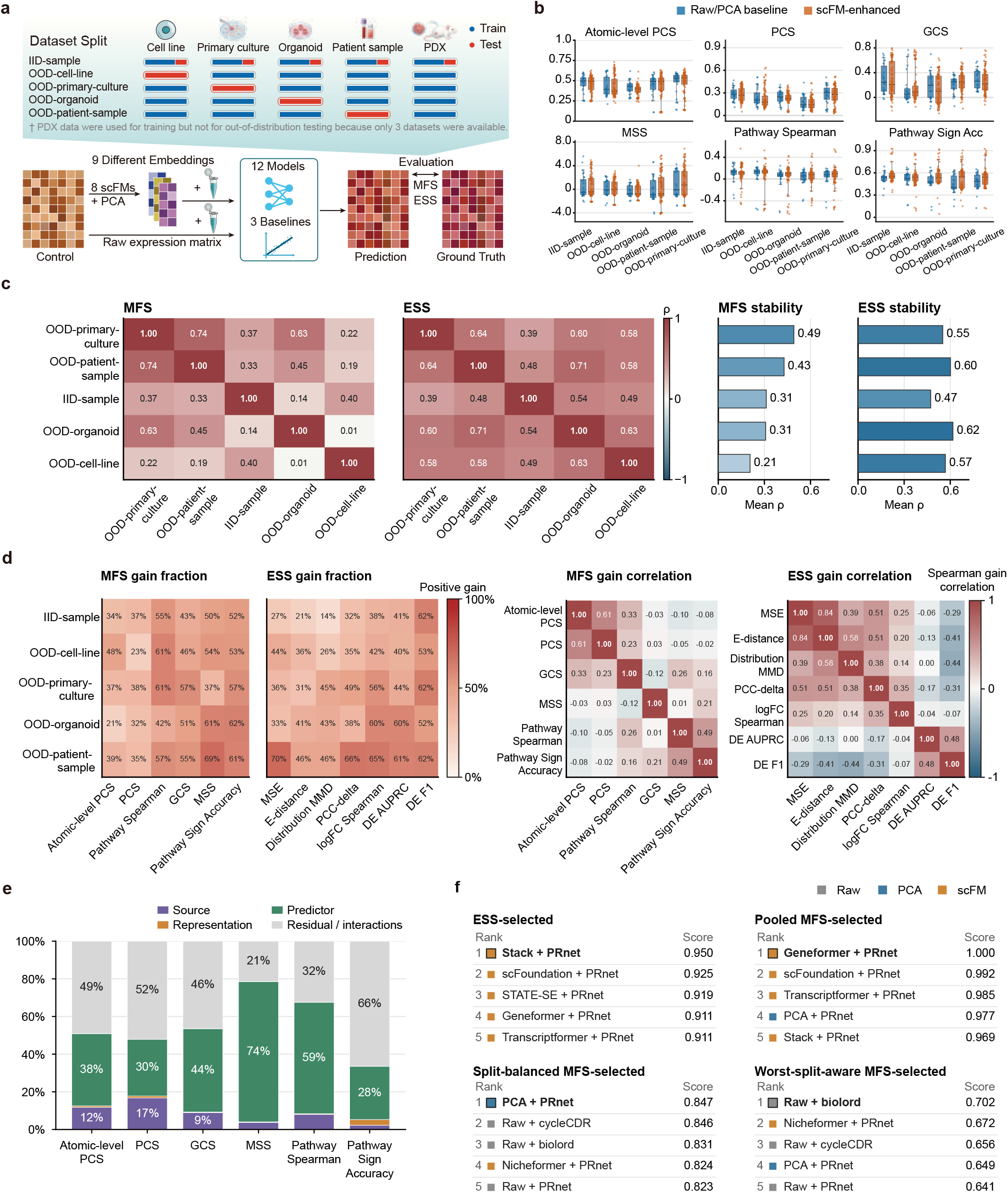
Source context reshapes mechanism fidelity and pipeline selection in multi-source perturbation benchmarks. **a**, Source-integrated benchmarking design across five source-aware splits. Raw expression input, PCA embeddings and frozen scFM embeddings were coupled to perturbation-prediction models and evaluated by MFS and ESS. PDX data were used only for training because this source contained too few datasets for out-of-distribution testing. **b**, Pipeline-level distributions of six mechanism-fidelity components across source-aware splits. Raw/PCA baseline pipelines and scFM-enhanced pipelines are compared for Atomic-level PCS, PCS, GCS, MSS, Pathway Spearman and Pathway Sign Accuracy. **c**, Cross-split stability of pipeline rankings induced by PCS-prioritized MFS and ESS. Heatmaps show pairwise Spearman correlations between matched pipeline rankings across source-aware splits, and adjacent bars show the mean rank concordance of each split with all other splits. **d**, Source- and metric-specific gains from frozen scFM embeddings relative to matched Raw-input predictors. Left heatmaps show the fraction of matched comparisons with positive gain for each MFS or ESS metric, and right heatmaps show Spearman correlations among metric-specific gains. **e**, Type II ANOVA variance partitioning of percentile-normalized mechanism-fidelity metrics in the balanced source–representation–predictor design. Stacked bars show variance fractions attributed to source, representation, predictor and residual or interaction effects. **f**, Top-five pipelines selected by four evaluation strategies: ESS, pooled PCS-prioritized MFS, split-balanced MFS and worst-split-aware MFS. Scores are rank-percentile summaries within each strategy, and colours denote Raw, PCA or scFM input types.

Source-aware splits showed distinct mechanism-fidelity profiles rather than a uniform shift in task difficulty (Fig. 5b). For each split, we summarized six mechanism-fidelity components across Raw/PCA baseline pipelines and scFM-enhanced pipelines. OOD-primary-culture retained relatively high Atomic-level PCS and PCS, whereas OOD-patient-sample showed weaker PCS, Pathway Spearman and Pathway Sign Accuracy despite Atomic-level PCS values close to iid-sample. OOD-cell-line was characterized mainly by lower PCS and GCS, whereas pathway-level metrics did not decrease uniformly. MSS also showed strong source dependence and broad pipeline heterogeneity. scFM-enhanced pipelines overlapped substantially with Raw/PCA baselines and did not produce a consistent upward shift across mechanism metrics. Thus, source context changed the mechanism-fidelity profile across gene-level, gene-set and pathway-level dimensions.

Source context also reordered pipeline rankings, particularly for mechanism fidelity (Fig. 5c). We compared the Spearman correlations of pipeline rankings induced by PCS-prioritized MFS and by ESS across the five source-aware splits. If source context only changed overall task difficulty, pipeline rankings would be expected to remain concordant across splits. Instead, MFS ranking correlations were low for several split pairs, ranging from 0.01 to 0.74. OOD-cell-line was the least concordant split, with a mean MFS rank correlation of 0.21 and a near-zero correlation with OOD-organoid (*ρ* = 0.01). By contrast, ESS rankings were more stable across splits, with pairwise correlations of 0.39–0.71 and mean correlations of 0.47–0.62. These results indicate that source context changes the relative mechanism-fidelity advantage of pipelines, not merely their absolute performance, and that this effect is stronger for mechanism fidelity than for expression reconstruction.

Matched Raw-input comparisons showed that scFM gains were source dependent and metric specific (Fig. 5d). Each scFM-enhanced pipeline was paired with the Raw-input pipeline using the same predictor and source-aware split, yielding 104 matched comparisons per split and 520 comparisons across splits. For MFS components, positive scFM gain was uncommon for core key-gene metrics: Atomic-level PCS improved in 21–48% of matched comparisons and PCS in 23–38%. Positive gains were more frequent for GCS and Pathway Sign Accuracy, but varied across sources and did not imply coordinated improvement in other mechanism dimensions. Gain correlations confirmed this dimensional specificity. Atomic-level PCS and PCS gains were moderately correlated (*ρ* = 0.61), whereas PCS gains were weakly related to GCS (*ρ* = 0.23), MSS (*ρ* = 0.03), Pathway Spearman (*ρ* = −0.05) and Pathway Sign Accuracy (*ρ* = −0.02). ESS gains were also heterogeneous: DE F1 improved in more than half of comparisons across all splits, whereas global distribution metrics often improved less frequently and sometimes showed negative gain correlations with DE F1. Thus, frozen scFM embeddings redistributed performance across sources and evaluation dimensions rather than producing a uniform improvement in perturbation prediction.

Variance partitioning further showed that representation identity was not the dominant determinant of performance (Fig. 5e). We percentile-normalized each metric across the balanced source–representation–predictor design and fitted an additive Type II ANOVA model, *Y* ∼ source + representation + predictor. For mechanism metrics, predictor choice accounted for the largest attributable main effect across all six components, ranging from 28.3% for Pathway Sign Accuracy to 74.5% for MSS. Source contributed more than representation for most mechanism metrics, including Atomic-level PCS (11.8% versus 0.7%), PCS (16.7% versus 1.0%), GCS (9.0% versus 0.2%) and Pathway Spearman (8.0% versus 0.4%). Representation accounted for only 0.2–3.0% of attributable variation. These results indicate that mechanism fidelity in the source-integrated regime was shaped more strongly by predictor choice and source context than by representation main effects alone. The corresponding ESS variance partitioning is shown in Supplementary Fig. S7.

The dependence on evaluation objective altered the preferred pipeline (Fig. 5f). ESS-based selection ranked Stack–PRnet first, and pooled PCS-prioritized MFS ranked Geneformer–PRnet first. Both strategies favoured scFM–PRnet combinations, but their winners had moderate weakest-split MFS values of 0.504 and 0.557, respectively. In contrast, split-balanced MFS selected PCA–PRnet, with a weakest-split MFS of 0.649, and worst-split-aware MFS selected Raw–biolord, with a weakest-split MFS of 0.702. The top-five sets also changed substantially: ESS and pooled MFS were dominated by scFM–PRnet combinations, whereas split-balanced and worst-split-aware selection included more Raw and PCA pipelines. Thus, aggregate or expression-based selection can prioritize pipelines with lower weakest-source mechanism fidelity, whereas source-aware MFS selection changes both the identity and input type of the preferred model.

Together, the source-integrated analysis shows that multi-source perturbation benchmarks require more than a single pooled performance summary. Source context reshaped mechanism-fidelity profiles, reordered MFS rankings, modulated scFM gains and contributed more variation than representation identity for most mechanism metrics. These results reinforce the need for mechanism-aware, source-stratified and source-balanced evaluation before interpreting a pipeline as source-general, and motivated failure-mode analysis of weak and source-dependent mechanism fidelity.

### 2.6. Hard-negative tests expose shortcut and failure modes in drug-response signature preservation

The weak and unstable mechanism fidelity observed above could arise from a trivial failure mode, in which models predict little or no perturbation, or from more specific shortcut behaviours. Hard-negative analyses supported the latter explanation. Current single-cell drug perturbation models usually generated non-zero transcriptional responses, but these responses were often not specific to the annotated drug-response signature. We therefore used hard-negative tests as signature stress tests, asking whether each prediction preserved the annotated response when compared with alternatives matched for expression background, gene identity, direction structure, generic response structure or global perturbation strength. These tests matched one nuisance property while disrupting the annotated mechanism, including the unperturbed state, the average training response, key-gene direction, effect magnitude, generic perturbation structure, non-matching signatures or global response strength. Across the 12 perturbation-prediction models and 3 baselines, the overall shortcut and failure burden ranged from 44.6% for PRnet to 61.5% for MAP (Fig. 6a). No-change collapse was 0% for all evaluated methods, indicating that the dominant failure was not absence of a perturbation signal. Instead, signature-specificity failure was pervasive, ranging from 84.3% for PRnet to 99.9% for MAP, and generic-response shortcut was also common, ranging from 54.3% for Squidiff to 88.9% for MAP. Lower-burden methods such as PRnet, cycleCDR, STATE-ST and baseMean still retained major hard-negative failures, whereas MAP, PrePR-CT, biolord, Squidiff and MLP showed broader shortcut profiles. Thus, limited mechanism fidelity arose from multiple shortcut and failure modes rather than from a single prediction error.

**Figure 6.**
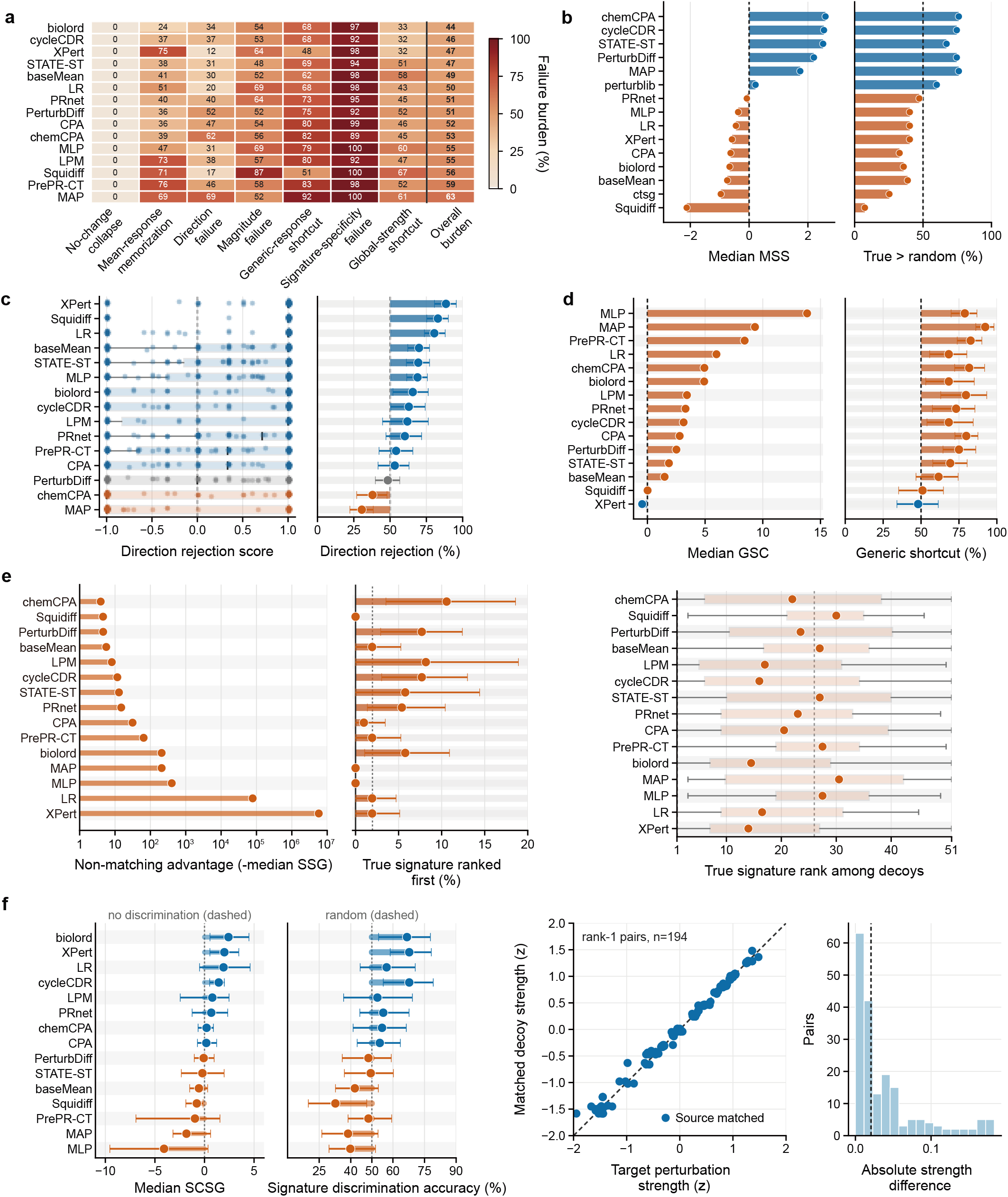
Hard-negative analyses reveal shortcut and failure modes in single-cell drug perturbation prediction. **a**, Model-level hard-negative failure spectrum. Heatmap values show the percentage of evaluable cases failing each of seven tests: no-change collapse, prediction close to the unperturbed control; mean-response memorization, prediction close to an average training response; direction failure, incorrect key-gene response polarity; magnitude failure, under- or over-estimated key-gene effect size; generic-response shortcut, stronger support for common perturbation signatures; signature-specificity failure, stronger support for non-matching drug-response signatures; and global-strength shortcut, reliance on perturbation magnitude rather than the case-specific signature. The rightmost column shows the overall shortcut and failure burden. **b**, Model-level enrichment of true key-gene responses over expression-matched random genes, showing the median MSS (left) and the fraction of cases in which true key genes scored higher than matched random genes (right). **c**, Directional consistency of key-gene responses, comparing annotated key-gene directions with sign-flipped signatures; subplots show the case-level direction rejection score distribution (left) and the true-direction preference rate with bootstrap confidence intervals (right). **d**, Generic-response shortcut burden, comparing true key-gene signatures with generic drug-response signatures; subplots show the median generic shortcut score (left) and the fraction of cases in which the strongest generic signature exceeded the true signature (right). **e**, Failure to prioritize true case-specific signatures over non-matching decoy signatures, summarized by non-matching advantage (left), the fraction of cases in which the true signature ranked first (middle) and the distribution of true-signature ranks among decoys (right). **f**, Strength-controlled discrimination between true signatures and response-strength-matched decoy signatures, showing the strength-controlled signature gap (first), discrimination accuracy (second), target–decoy strength matching (third) and absolute strength differences after matching (fourth).

We first tested whether predicted effects were enriched in literature-curated key genes rather than expression-matched random genes (Fig. 6b and Supplementary Fig. S8a). The MSS compared predicted support for true key genes with support for random genes matched by expression properties. Key-gene specificity was strongly model dependent. PRnet showed the strongest enrichment, with a median true-versus-random advantage of 5.17 and true key genes scoring above random genes in 81.9% of cases; cycleCDR and PerturbDiff also showed positive advantages of 3.32 and 3.08. However, the median MSS across all 15 evaluated methods was -0.47, and Squidiff showed the weakest enrichment, with a median advantage of -2.61 and a true-greater-than-random fraction of 5.9%. Because the negative genes were expression matched, these failures reflect loss of signature localization rather than trivial differences in gene detectability.

Direction preservation was a separable and partially easier task, but it remained insufficient for full mechanism fidelity (Fig. 6c). We constructed a sign-flipped hard negative by retaining the same key genes but reversing their annotated up- or down-regulation direction. XPert, Squidiff and linear regression showed the strongest true-direction preference rates, 85.3%, 83.5% and 79.0%, respectively. However, PrePR-CT was close to random, with a true-direction preference rate of 54.1%, and PerturbDiff and MAP showed negative median direction advantages. Thus, some models preserved response polarity, but correct direction alone did not ensure preservation of the correct case-specific signature.

We next tested whether predictions favoured generic drug-response signatures over the current case-specific signature (Fig. 6d and Supplementary Fig. S8b). This comparison targeted a common shortcut: reproducing a recurrent perturbation programme rather than the signature for the current drug, dose, time and cellular context. All models had a positive median generic-response shortcut score. MAP showed the strongest shortcut, with a median score of 10.32 and generic signatures exceeding true signatures in 88.9% of cases; PrePR-CT and biolord also showed large median generic advantages, 8.43 and 7.37. Even the least affected models retained substantial shortcut behaviour, with generic-over-true fractions of 54.3% for Squidiff, 61.1% for XPert and 61.5% for baseMean. These results show that many predictions resembled plausible perturbation responses but were generic rather than case specific.

The strongest evidence for limited case-specific signature fidelity came from non-matching signature decoys (Fig. 6e and Supplementary Fig. S9a). For each case, we compared the true key-gene signature with signatures from other perturbation cases with different drugs and low key-gene overlap. No model had a positive median signature-specificity gap, meaning that the strongest non-matching signature usually received higher support than the true signature. Even the relatively better methods had negative gaps, including Squidiff (-4.21), baseMean (-5.64) and PRnet (-6.34). PRnet ranked the true signature first most often, but only in 14.9% of cases, and the median true-first fraction across all evaluated methods was about 1.9%. Thus, model predictions frequently supported a biologically plausible but incorrect signature more strongly than the annotated signature for the current case.

Finally, we controlled for global perturbation strength by matching each target case to a decoy with similar response magnitude, different drug identity and minimal key-gene overlap (Fig. 6f and Supplementary Fig. S9b). The matched negative design placed most target–decoy pairs near the equal-strength diagonal, focusing the comparison on drug-specific signatures rather than response magnitude. Only a subset of models retained stable discrimination after this control. PRnet performed best, with a median strength-controlled signature gap of 2.82 and a discrimination accuracy of 72.5%; linear regression and cycleCDR also showed positive gaps, 2.12 and 1.50. By contrast, the median discrimination accuracy across all evaluated methods was approximately 48.7%, and the median gap was -0.21. These results indicate that global response strength can itself act as a shortcut: a model may match how strong a perturbation appears while missing the key-gene signature that distinguishes it from another response of similar magnitude.

Together, these analyses show that hard negatives are necessary for separating signature fidelity from non-specific shortcut and failure modes. Current models rarely collapsed to the control state, but they frequently misplaced signal onto expression-matched random genes, confused response direction, favoured generic signatures, prioritized non-matching case signatures or relied on global perturbation strength. Hard-negative testing therefore complements expression similarity metrics and aggregate mechanism fidelity scores by exposing when an apparently plausible perturbation response does not preserve the annotated drug-response signature.

## 3. Discussion

scDrugPerturb-Bench shows that preserving the transcriptome is not the same as preserving the curated drug-response programme. Across direct prediction, representation-augmented prediction, source generalization and downstream retrieval, models often produced plausible non-zero responses, but those responses did not reliably preserve key-gene direction, pathway polarity or mechanism specificity. The benchmark therefore makes mechanism fidelity visible as a separate axis of model quality, rather than a guaranteed consequence of expression similarity.

This separation changes how perturbation models should be evaluated. Conventional expression-similarity metrics remain useful for reconstruction, correlation and differential-expression agreement, but they were weak proxies for signature-level fidelity. ESS often selected different top pipelines from PCS-prioritized MFS, and the downstream transcriptome-based drug-retrieval analysis showed that MFS can provide a more informative signal for drug retrieval. The implication is simple: expression-level scores are appropriate for descriptive response modelling, whereas mechanism-guided use cases require direct assessment of key-gene direction, effect size, pathway response and mechanism specificity.

The results also caution against equating larger models or pretrained representations with better biological fidelity. More complex perturbation architectures did not consistently outperform simpler baselines, and their advantages depended on the split, the predictor and the metric. Frozen scFM embeddings could help in selected settings, but the gains were local and did not generalize across all mechanism dimensions. Practical deployment adds another constraint, because runtime, RAM and GPU memory varied substantially across methods. Together, these observations suggest that architecture scale is insufficient without objectives that explicitly localize signal to the annotated mechanism.

Source context emerged as a major source of instability. In the multi-source benchmark, source composition changed absolute performance, reordered pipeline rankings and altered the identity of the preferred representation. Pooled summaries therefore conceal an important boundary: a pipeline that looks strong on average may be unstable, or even poor, in a specific source. For benchmarks meant to support cell lines, primary cultures, organoids and patient samples, source-stratified and source-balanced reporting is essential before any model is described as broadly generalizable.

The hard-negative analyses help explain these failures. Most models did not collapse to a no-change prediction, but many reused generic response programmes, misplaced signal onto non-specific genes, or relied on perturbation strength instead of the case-specific signature. These shortcuts can produce plausible profiles while still assigning stronger support to a decoy than to the annotated response. Hard negatives therefore act as signature stress tests, and they are valuable both as evaluation criteria and as training constraints.

Several limitations define the scope of these conclusions. scDrugPerturb-Bench relies on literature-curated mechanism annotations, so its labels are directional summaries rather than complete mechanistic descriptions. The benchmark also evaluates transcriptional mechanism fidelity, not target engagement, protein activity or clinical benefit. MFS further depends on the chosen mechanism metrics and their aggregation, so alternative weightings or additional readouts could change rankings in specific applications. scFM embeddings were evaluated as frozen representations, so different outcomes may emerge with fine-tuning or architecture-specific adapters. Finally, coverage remains uneven across drugs, doses, times and cell states. Even so, the benchmark provides a practical framework for separating reconstruction from mechanism-aware prediction, and for testing whether a model is suitable for mechanism-guided retrieval.

## 4. Methods

### 4.1. Models

#### Drug perturbation prediction

We selected benchmark models to cover the main algorithmic families used for single-cell drug perturbation prediction, together with simple baselines that test whether more complex architectures outperform average-response or linear predictors. Each model was trained and evaluated using the same split-specific data partitions. Training used only the training set, and all reported metrics were computed on held-out test cases. Literature-curated mechanism annotations were used only for evaluation and were not provided to the models during training.

The direct perturbation-prediction benchmark included 12 published models and three baseline predictors. The baselines were linear regression (LR), an MLP and a mean-response predictor[25]. The published models comprised five VAE-based approaches: CPA[3], chemCPA[4], cycleCDR[26], PRnet[27] and biolord[28]. Four models used transformer architectures: STATE-ST[29], XPert[7], MAP[30] and LPM[8]. We also included two diffusion-based models, Squidiff[5] and PerturbDiff[6], and the graph neural network-based model PrePR-CT[9]. We used public implementations and pretrained components where available. All models received inputs derived from the same processed expression matrices and metadata. Method-specific parameters followed official documentation or released benchmark settings, with only format-level changes needed to align each tool with the common data splits.

We separately evaluated single-cell foundation models (scFMs) and PCA as upstream representation models. This analysis included scGPT[31], scFoundation[11], Geneformer[12], Transcriptformer[32], Nicheformer[33], SCimilarity[34], Stack[35] and STATE-SE[29], with PCA included as a non-neural representation baseline. For each scFM, processed cells were mapped to the model-specific gene space, embedded with released model weights and reduced to a single cell-level vector using the embedding layer and pooling rule prescribed by that model. These rules differed across models, including CLS-token extraction, mean pooling over gene tokens, flattened token grids and direct MLP-based cell encoders. Genes absent from a model vocabulary were discarded or zero-filled according to the corresponding implementation, and gene identifiers were aligned by exact gene-symbol matching or symbol-to-Ensembl mapping where required. PCA embeddings were computed as a non-neural baseline from highly-variable genes in the processed expression matrices. All embeddings were generated with model-specific preprocessing and tokenization rules, including expression binning, rank-value encoding, fixed canonical gene orders or position-based token assignment where specified by the released implementation. All embeddings were kept fixed and provided to each compatible downstream perturbation-prediction model under the same data splits. This embedding experiment therefore tested whether an upstream transcriptomic representation improved perturbation prediction across downstream model architectures rather than under a single predictor. PrePR-CT was excluded from this embedding experiment because its graph neural network architecture could not be directly connected to external cell-embedding matrices within the common interface. Model-specific embedding dimensions, layers, pooling operations and gene-alignment rules are provided in Supplementary Table S1.

#### Transcriptome-based drug retrieval

To assess whether MFS or ESS was more closely aligned with downstream drug-retrieval utility, we used CURE[23] as a fixed transcriptome-based drug-retrieval model. The same nine upstream representation methods, comprising eight scFMs and PCA, were evaluated through a common perturbation-prediction and retrieval pipeline so that differences in downstream performance could be attributed to representation selection rather than to the retrieval architecture. For each representation, the predicted transcriptomic response was supplied to CURE, which mapped the response to a query drug fingerprint. This fingerprint was compared with a reference library of 8,495 small-molecule fingerprints, and candidate compounds were ranked by fingerprint similarity. The top 500 ranked compounds were retained for retrieval evaluation using Hit@K, normalized discounted cumulative gain at *K* (NDCG@K) and mean reciprocal rank (MRR). This common downstream model enabled an application-level comparison of whether representations prioritized by MFS or ESS more effectively supported retrieval of the true perturbing drug.

### 4.2. Benchmark datasets and preprocessing

#### 4.2.1. Data collection and case curation

We constructed scDrugPerturb-Bench through a structured literature-to-data curation workflow. In April 2026, PubMed was searched for English-language articles published from 2020 to 2026 using the query `scRNA-seq[All Fields]` AND `2020:2026[dp]` AND `english[Language]` . The search returned 54,549 records. Titles and abstracts were screened with a recall-oriented DeepSeek-V4-Pro prompt at temperature 0.1 to identify human single-cell RNA-seq studies after pharmacological perturbation, with treated and matched-control expression profiles and publicly accessible data. Records with incomplete information were retained for full-text review. This stage yielded 4,228 articles for which we retrieved the full-text PDF.

Full-text screening and structured extraction used the main article PDF, including figures and figure captions, together with publicly available expression data. Supplementary materials were not used. Claude Code with DeepSeek-V4-Pro performed full-text assessment and extraction, and Codex with GPT-5.5 provided an independent automated check. Eligible datasets had to contain human single-cell transcriptomic profiles from a single small-molecule treatment, a matched untreated, vehicle or baseline control, publicly accessible expression data and at least one experimentally supported gene-level change with an explicit direction. We excluded non-human studies, records without drug-treated single-cell data or a matched control, macromolecular perturbagens, multi-drug treatments without an isolatable single-drug arm, drug-resistance-state comparisons without an acute treatment contrast and studies without an explicit directional gene-level claim. The final curation set comprised 93 studies, 181 datasets and 423 cases.

Eligible studies were represented using predefined dataset-level and case-level schemas. Dataset-level fields included publication and accession identifiers, tissue and cellular context, experimental platform, perturbagen, control, and dose and time design. Case-level fields included source location, verbatim source statement, drug and cell context, mechanism gene or gene set, reported direction, control, dose, treatment time and evidence type. Language models were used to locate candidate evidence and populate these fields, rather than to generate mechanism annotations. A gene and its direction were retained only when supported by an explicit statement, figure or figure caption in the source article; directions were not inferred from general biological knowledge. Each case was therefore linked to a verbatim source passage and its location in the original article.

All cases were assigned among five human curators, with one curator per case and no duplicate review. Curators checked cellular context, mechanism genes and directions, dose, treatment time and source passage against the original article. Manual review covered all retained cases. For all cases, we downloaded the deposited data, converted them into gene-expression matrices and performed cell annotation when required. We then tested whether treatment-control expression changes agreed with the extracted direction and performed an additional quality-control assessment of each processed matrix.

#### 4.2.2. Data preprocessing

##### Case filtering

Before model evaluation, we used Hedges’ *g* to quantify each treatment-control effect in the processed expression data and compared its sign with the literature-curated gene-direction annotation. If the observed effect matched the reported direction and its 95% confidence interval excluded zero, the gene was labelled UP or DOWN accordingly. If the point estimate matched the reported direction but the confidence interval included zero, the gene was labelled not significant (NS). Cases were excluded when the observed effect pointed in the opposite direction to the source conclusion. This retained directionally concordant but statistically inconclusive observations while removing cases that contradicted the source evidence.

##### Cell filtering

We applied a consistent cell-level quality-control workflow to all single-cell perturbation datasets, using raw count matrices whenever available. Cells were removed if they had fewer than 200 detected genes, fewer than 500 UMIs or a mitochondrial fraction above 50%. Among cells that passed these thresholds, potential doublets were identified with Scrublet using its automatically selected cutoff. We then performed sample-specific outlier filtering based on the median absolute deviation. Total UMIs and detected genes were evaluated in log space, whereas the fraction of reads assigned to the top 20 genes and the mitochondrial fraction were filtered only for high-end outliers. Cells that passed QC were retained for gene mapping and downstream filtering.

##### Gene filtering and mapping

Because the datasets came from multiple studies with heterogeneous gene annotations, we harmonized gene identifiers to current HGNC-approved symbols using GENCODE v44 and HGNC reference annotations[36, 37]. The mapping procedure accounted for current and deprecated symbols, Ensembl and HGNC identifiers, aliases and embedded identifiers, whereas ambiguous identifiers were not reassigned. After harmonization, we retained only genes in the predefined gene set, including regulatory-sequence genes, protein-coding genes and transcription factor genes. When multiple source features mapped to the same approved gene symbol, their expression values were summed to produce a single non-redundant gene feature.

##### Data normalization

We normalized the filtered count matrices with the same Scanpy-based global scaling strategy [38]. For each cell, counts were rescaled to 10,000 total counts and log-transformed. This reduced sequencing-depth variation while preserving expression differences more likely to reflect biological responses.

#### 4.2.3. Data splits

We evaluated perturbation prediction under two complementary generalization regimes. The cell-line regime included IID-sample, OOD-drug, OOD-cell, OOD-tissue and OOD-drug-cell-pair splits. These splits tested generalization across held-out perturbations, cellular contexts, tissues and specific drug–cell combinations. The cross-source regime assessed transfer from one set of experimental sources to held-out source domains. Splits were constructed by dataset. Matched control profiles, perturbed profiles and case annotations from the same dataset were assigned to a single training or test partition. Detailed split construction is provided in the Supplementary Methods.

### 4.3. Evaluation

All metrics were computed on held-out test cases. A test case *i* specifies a perturbation condition *d*_*i*_, including its drug and dose. The set *G*_*i*_ contains the literature-curated mechanism genes, whereas *G*_*i*_ denotes the aligned expression gene set used for transcriptome-wide similarity metrics. The model index is omitted from case-level metric definitions for readability.

#### 4.3.1. Mechanism-fidelity score

Mechanism fidelity was assessed using six complementary metrics. Hedges’ *g* provided the common effect-size basis for the gene-level mechanism metrics and was not treated as an additional MFS component.

For a gene–dose unit (*g, d*) and data source *q* ∈ {real, pred}, let 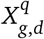 denote its cell-level expression values under condition *d*. The corresponding matched-control values are denoted 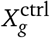. We quantified the perturbation effect as Hedges’ *g*, a small-sample-corrected standardized mean difference:

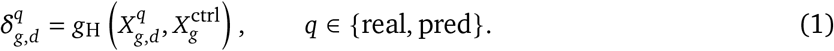

Positive and negative values indicate upregulation and downregulation relative to control, respectively. Bootstrap confidence intervals used 2,000 cell-level resamples, with perturbed and matched-control cells resampled separately. The pooled variance, correction factor and full bootstrap procedure are defined in Supplementary Methods.

##### Pattern Consistency Score (PCS)

PCS measures whether predicted perturbation effects preserve the observed direction of annotated gene responses. Predictions were considered confident when the bootstrap 95% confidence interval of 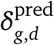 did not include zero. At the atomic level, PCS is a gene-direction accuracy indicator for each confident annotated gene–dose unit:

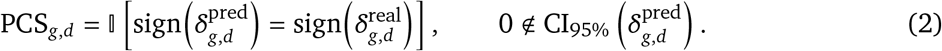

Units with confidence intervals that included zero were treated as inconclusive. For the confident subset C_*i*_ of annotated gene–dose units in case *i*, case-level PCS used a strict consistency rule:

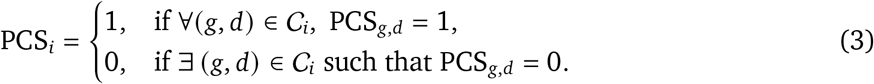

Cases with |C_*i*_| = 0 were unevaluable and excluded from PCS aggregation.

##### Effect Size Recovery (ESR)

ESR measures calibration of perturbation-effect magnitude after directional consistency has been established. For the set E_*i*_ of confident, directionally correct annotated gene–dose units with sufficiently large observed effects 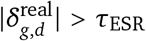, with *τ*_ESR_ = 0.2), case-level ESR was the median predicted-to-observed effect-size ratio:

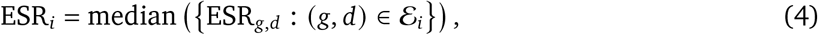

where the ideal value is 1.

##### Gene-set Coherence Score (GCS)

GCS measures whether a model preserved coordinated responses within an annotated gene set. For each test case *i*, predicted and observed Hedges’ *g* values over_*i*_ were arranged into vectors 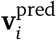 and 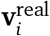. GCS was defined as their Spearman rank correlation:

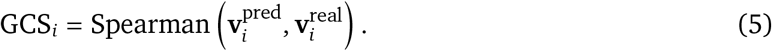

GCS was treated as undefined when the gene set was too small or either effect-size vector was constant.

##### Mechanism Specificity Score (MSS)

MSS tests whether predicted effects are enriched in mechanism-relevant genes rather than distributed across background genes. For each test case, the mean absolute predicted effect over *G*_*i*_, denoted 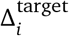, was compared with a null distribution from *R* = 1,000 size-matched random gene sets, denoted 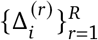. MSS was defined as a robust standardized contrast:

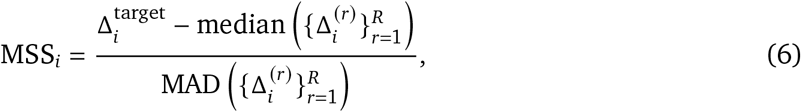

with zero-MAD null distributions handled as described in Supplementary Methods. Positive MSS values indicate stronger predicted effects in the annotated target genes than in matched random gene sets.

For each case, genes were ranked by their log fold changes relative to matched control, computed with a numerical-stability constant *ϵ* = 10^−6^. Predicted and observed profiles were then analysed separately by pre-ranked gene-set enrichment analysis using KEGG 2021 Human pathway gene sets. For pathway *k*, 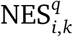, where *q* ∈ {real, pred}, denotes the enrichment score normalized against 1,000 size-matched random gene sets. Positive and negative NES values indicate enrichment among upregulated and downregulated genes, respectively.

##### Pathway Spearman

Pathway Spearman evaluates concordance between the predicted and observed NES profiles:

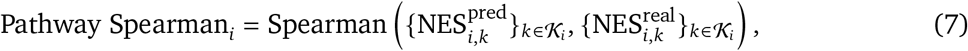

where *K*_*i*_ contains pathways with finite predicted and observed NES values.

##### Pathway Sign Accuracy

Pathway Sign Accuracy measures agreement in pathway polarity among observed perturbed pathways:

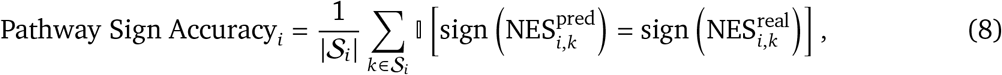

where *S*_*i*_ contains pathways with observed NES magnitude above *τ*_pathway_ = 1.0. Pathway metrics were undefined when their required pathway sets were empty or too small for correlation analysis. Full implementation details, thresholds and undefined-case handling are provided in Supplementary Methods.

#### 4.3.2. Expression-similarity score

Conventional transcriptomic similarity was evaluated using seven metrics over the aligned gene set *G*_*i*_, which is distinct from the mechanism gene set *G*_*i*_. These metrics used three case-level quantities: mean expression under the predicted or observed perturbed condition, mean perturbation effects relative to matched control and log fold changes relative to matched control.

We organized the seven metrics into three complementary groups. Distribution-level agreement was measured by Mean Squared Error (MSE), which quantifies the average squared discrepancy between predicted and observed mean perturbation-effect vectors; E-distance, which contrasts cross-population distances with within-population distances between predicted and observed cells; and Distribution MMD, which compares predicted and observed cell populations using an RBF-kernel maximum mean discrepancy with the kernel bandwidth selected by the median pairwise-distance heuristic. Perturbation-effect agreement was measured by PCC-delta, the Pearson correlation between predicted and observed mean perturbation-effect vectors, and logFC Spearman, the Spearman correlation between gene-level log fold-change rankings. Differential-expression recovery was evaluated by DE AUPRC, which treats observed differential-expression calls as labels and absolute predicted log fold changes as scores, and DE F1, which compares observed and predicted binary differential-expression calls at the same absolute log2 fold-change threshold (*τ*_DE_ = 0.25). Lower MSE, E-distance and Distribution MMD values indicate better agreement. Log fold changes used*ϵ*= 10^−6^. Full metric formulas, thresholds, kernel bandwidth selection and undefined-case handling are provided in Supplementary Methods.

#### 4.3.3. Drug-retrieval evaluation

CURE was used as a fixed transcriptomic retrieval model because it accepts predicted transcriptomic profiles as queries for drug retrieval. It was not treated as a perturbation-prediction baseline. Instead, CURE provided a common downstream test of whether pipelines selected by MFS or ESS improved retrieval of the true drug. The reference library contained 8,495 small molecules. It was constructed from DrugBank 5.1.22 by retaining 8,486 DrugBank compounds with available structures and identifiers up to DB13920, corresponding to 64.5% DrugBank coverage. Approximately 4,700 higher-identifier DrugBank entries, mostly experimental compounds without public SMILES or PubChem CID annotations, were not included. Among the 48 benchmark cell-line compounds, 39 overlapped with the DrugBank fingerprint library and 9 additional benchmark compounds were added, yielding the final 8,495-compound library. Compounds were represented using Morgan fingerprints (ECFP4; radius = 2; 2,048-bit count vectors), and candidate ranking used the Tanimoto, or Jaccard, similarity coefficient. Predicted perturbation profiles from the selected pipelines were supplied to CURE and evaluated using Hit@K, NDCG@K and mean reciprocal rank (MRR). The composite early and full-range Drug Retrieval Scores, together with their ranking rules, are defined in the Methods ranking section below.

### 4.4. Methods ranking

We ranked models separately for mechanism fidelity, expression similarity and downstream drug retrieval, because these scores measure different evaluation targets and were not intended to be collapsed into a single global objective. All composite scores were computed after the corresponding case-level metrics had been aggregated to the dataset–model level. Unless otherwise specified, ranks were assigned within each dataset, with rank 1 denoting the best-performing model.

For any dataset–model pair *u* = (*j, m*) and metric *S*, values were first oriented so that larger values indicated better performance and then robustly standardized:

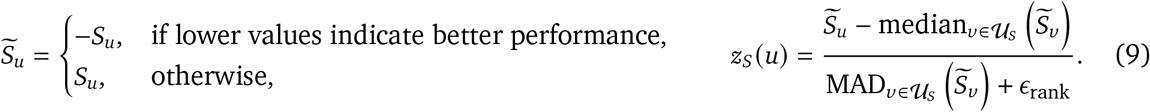

where *:ϵ*_rank_ is a small stabilizer used only for robust rank standardization. Median aggregation was used throughout the composite scores to reduce sensitivity to individual metrics with extreme values or nested *K*-dependent definitions.

#### Mechanism-fidelity ranking

Mechanism-fidelity ranking used a PCS-prioritized MFS rule. For each dataset–model pair (*j, m*), dataset-level PCS was calculated as a weighted mean over valid test cases,

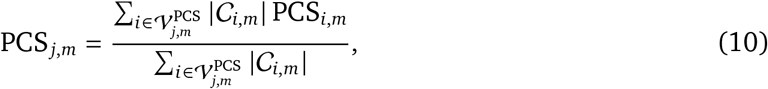

so that cases with more confident mechanism genes contributed proportionally more information. Each remaining mechanism metric was summarized by the median across its valid cases. PCS was used as the primary ordering variable because it directly measures whether the predicted response preserves the annotated direction of key-gene effects. Models within *δ*_PCS_ = 0.02 of a tier anchor were assigned to the same PCS tier. Within each tier, models were ranked by an auxiliary mechanism score:

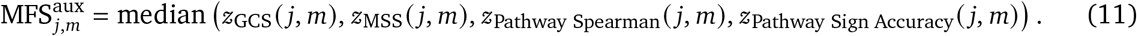

The final MFS ranking sorted models by ascending PCS tier, descending MFS^aux^ and descending PCS. This rule allows GCS, MSS and pathway-level metrics to refine ranking among models with similar directional key-gene consistency, while preventing secondary metrics from overriding a substantial PCS difference. ESR was not included in the primary MFS ranking because it is a ratio-based calibration metric computed only after confident directional consistency. We therefore reported ESR separately and used it for sensitivity analyses of effect-size calibration. PCS, ESR, GCS, MSS, Pathway Spearman and Pathway Sign Accuracy were also reported separately.

#### Expression-similarity ranking

Expression-similarity ranking used an ESS that combined seven conventional transcriptomic similarity metrics. For each metric, the dataset–model score was the median across valid test cases. Error and distance metrics were sign-oriented before standardization so that larger *z*-scores consistently indicated better performance. The seven metrics were grouped into global distributional, perturbation-effect and differential-expression components:

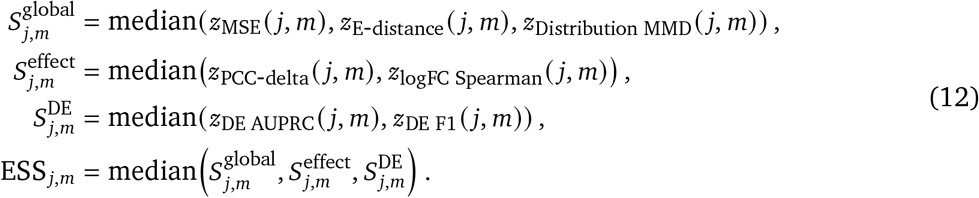

Expression-similarity ranks were assigned by sorting ESS in descending order within each dataset. Cases for which a metric was undefined were excluded only from the aggregation of that metric. Pathway Spearman and Pathway Sign Accuracy were excluded from ESS and used only for mechanism-fidelity ranking.

#### Drug-retrieval ranking

Drug-retrieval ranking summarized downstream retrieval performance while emphasizing early enrichment of the true drug among candidate compounds. We grouped nested *K*-dependent Hit@K and NDCG@K metrics to avoid over-weighting repeated cutoffs. Early summaries used *K* = {5, 10, 20, 50}, whereas broad summaries used *K* = {100, 500}:

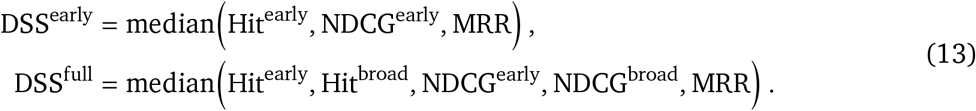

Mean reciprocal rank (MRR) was retained as a separate first-hit component. In the drug-retrieval analysis, representations were ranked by descending DSS^early^, and DSS^full^ was used as a sensitivity analysis. Detailed grouping and aggregation rules are provided in Supplementary Methods.

#### Integrated representation-selection score

For the sensitivity analysis in Fig. 3d, we defined an integrated representation-selection score to combine mechanism fidelity and expression similarity during representation selection. In each OOD split *s*, representations *e* were ranked separately by PCS-prioritized MFS Overall and ESS Overall, and ranks were converted to direction-oriented percentiles *P*_MFS_(*s, e*) and *P*_ESS_(*s, e*), with larger values indicating better performance. The integrated score was

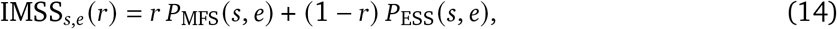

where *r* ∈ {0.1, 0.2, …, 1.0} controlled the weight assigned to mechanism fidelity. For each *r* and split, the representation with the largest IMSS was selected and evaluated using the downstream CURE-based drug-retrieval metrics. Relative changes in Hit@K and NDCG@K were computed against the ESS-selected representation, and 95% confidence intervals were estimated with 5,000 benchmark-level bootstrap resamples.

### 4.5. Hard-negative analysis

Hard-negative analysis was designed to identify shortcut solutions that can yield apparently plausible perturbation predictions without preserving the case-specific drug-response signature. Each evaluable unit was a test case *i*, defined by study, cell context, drug, dose and time, and each prediction was indexed by model *m*. These analyses used pseudobulk perturbation-effect vectors for case-level tests and predicted Hedges’ *g* values for gene-level mechanism tests.

We prespecified seven failure modes. No-change collapse tested whether the predicted response norm was close to the unperturbed state. Mean-response memorization tested whether predictions were closer to a training-set average response than to the corresponding observed response. Key-gene specificity tested whether predicted effects were stronger in literature-curated key genes than in expression-matched random genes. Direction-flipped decoys tested whether predicted effects supported the annotated directions more strongly than the same genes with reversed signs. Magnitude failure tested whether key-gene effect sizes were substantially under- or over-estimated. Generic-response shortcuts tested whether predictions supported training-derived generic drug-response signatures more strongly than the case-specific signature. Non-matching and strength-matched signature decoys tested whether the prediction preferred other curated signatures or signatures with similar global perturbation strength.

For each failure mode *k*, we defined a binary indicator *F*_*i,m,k*_ when the corresponding hard-negative criterion was evaluable. The model-level burden for mode *k* and the overall shortcut burden were

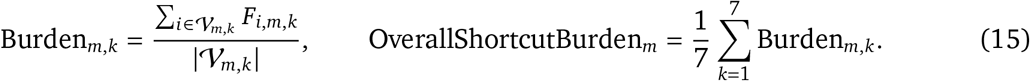

Here, *V*_*m,k*_ denotes the cases evaluable for model *m* and failure mode *k*. The resulting burden profile summarizes whether poor mechanism fidelity arises primarily from collapse, memorization, direction reversal, magnitude miscalibration, generic-response shortcuts, loss of signature specificity or reliance on global perturbation strength. Detailed hard-negative definitions and thresholds are provided in Supplementary Methods.

### 4.6. Data availability

The processed scDrugPerturb-Bench data are available from Hugging Face at https://huggingface.co/datasets/mindflow-cn/scDrugPerturb-Bench.

### 4.7. Code availability

The source code for scDrugPerturb-Bench benchmark construction, model evaluation and analysis is available on GitHub at https://github.com/mindflow-cn/scDrugPerturb-Bench. The code and skills used for literature-based dataset collection are available at https://github.com/mindflow-cn/paper2perturb. The project website is available at https://mindflow-cn.github.io/simucella/paper.html.

---

**Figure S1.**
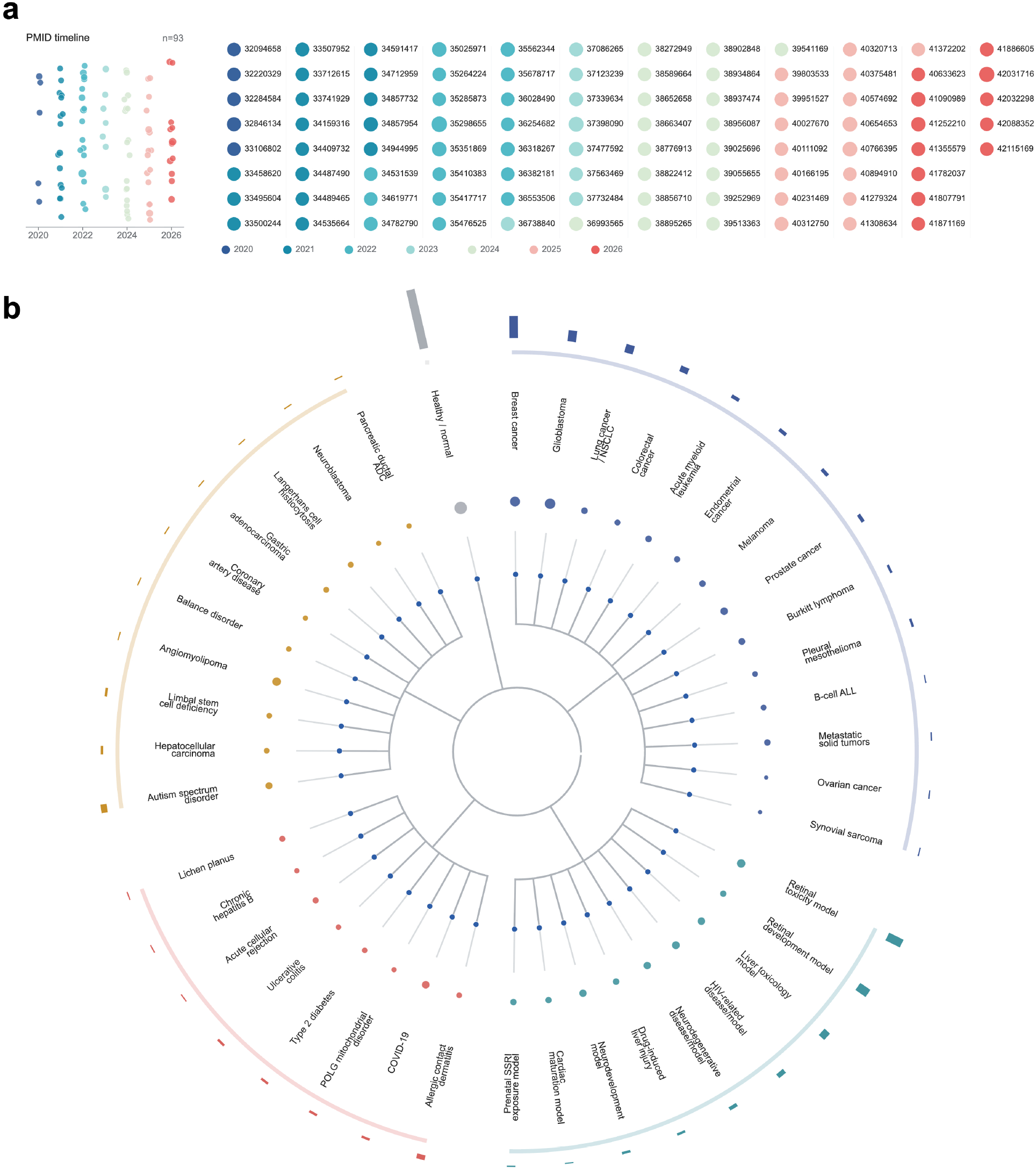
Publication timeline and disease-state coverage of the curated benchmark. **a**, PMID-level distribution of the source literature used for data collection, spanning 2020 to April 2026. Circle size denotes the number of datasets contributed by each publication. **b**, Coverage of 42 harmonized disease or state categories represented in the collected datasets. The central radial hierarchy groups categories into broader disease or state classes, and each outward branch corresponds to one harmonized category labelled at the perimeter. Colours denote major classes, including healthy or normal states, tumour-related diseases, developmental or toxicology models, immune, infection and metabolic states, and other disease states. Outer bars indicate the number of datasets assigned to each category, whereas inner points indicate relative cell abundance. The distribution shows broad but uneven coverage, with healthy or normal states and tumour-related categories forming the largest components; breast cancer and glioblastoma have relatively high dataset or cell representation, and retinal toxicity or development, hepatotoxicity and HIV-related models provide additional non-tumour perturbation contexts.

**Figure S2.**
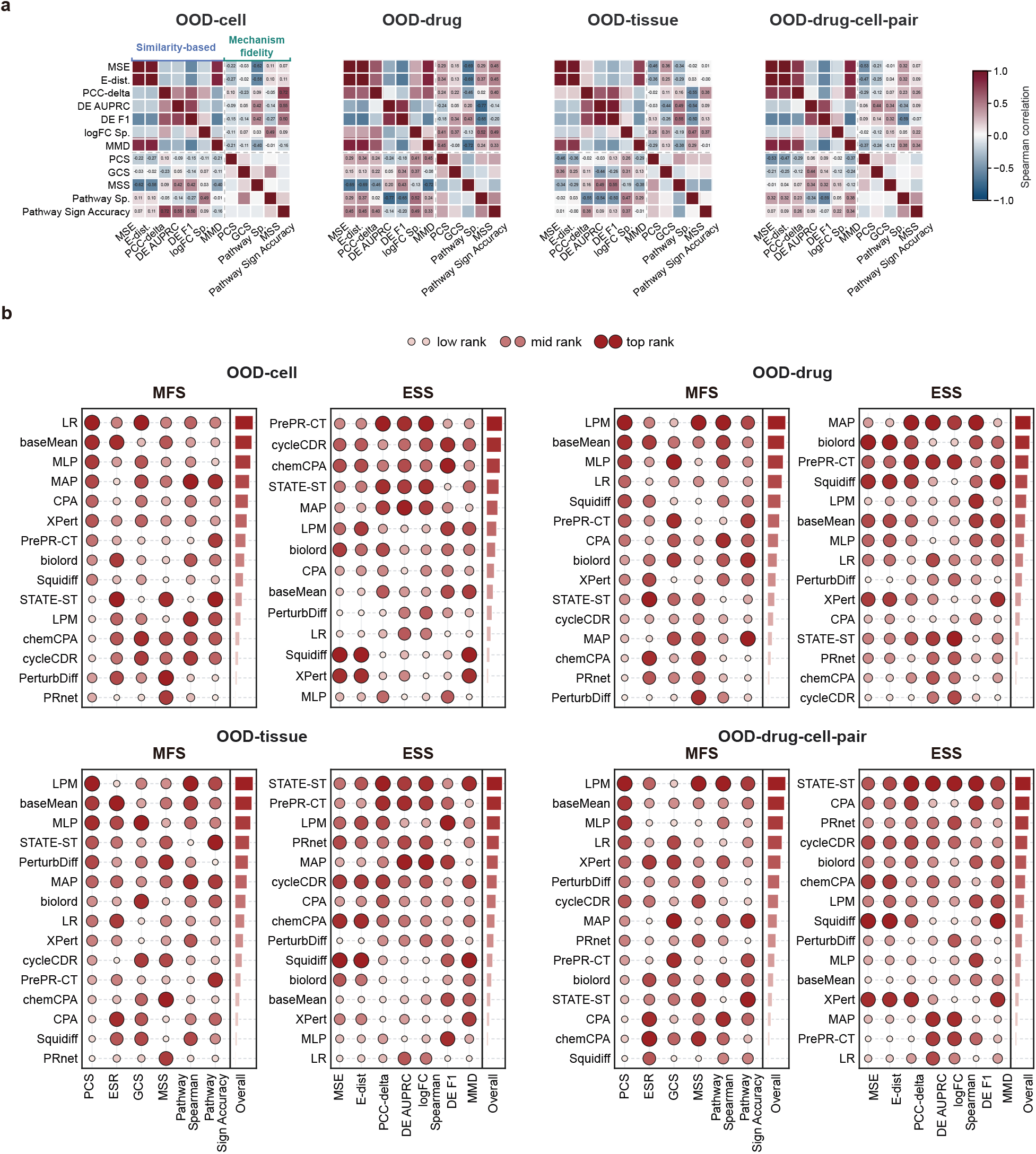
Supplementary analyses of expression-similarity and mechanism-fidelity discordance. **a**, OOD-specific Spearman correlation matrices corresponding to Fig. 2c, shown for OOD-cell, OOD-drug and OOD-tissue, OOD-drug-cell-pair settings. **b**, OOD-specific comparisons of model rankings induced by the Expression Similarity Score and the Mechanism Fidelity Score, corresponding to Fig. 2d, highlighting their rank discordance.

**Figure S3.**
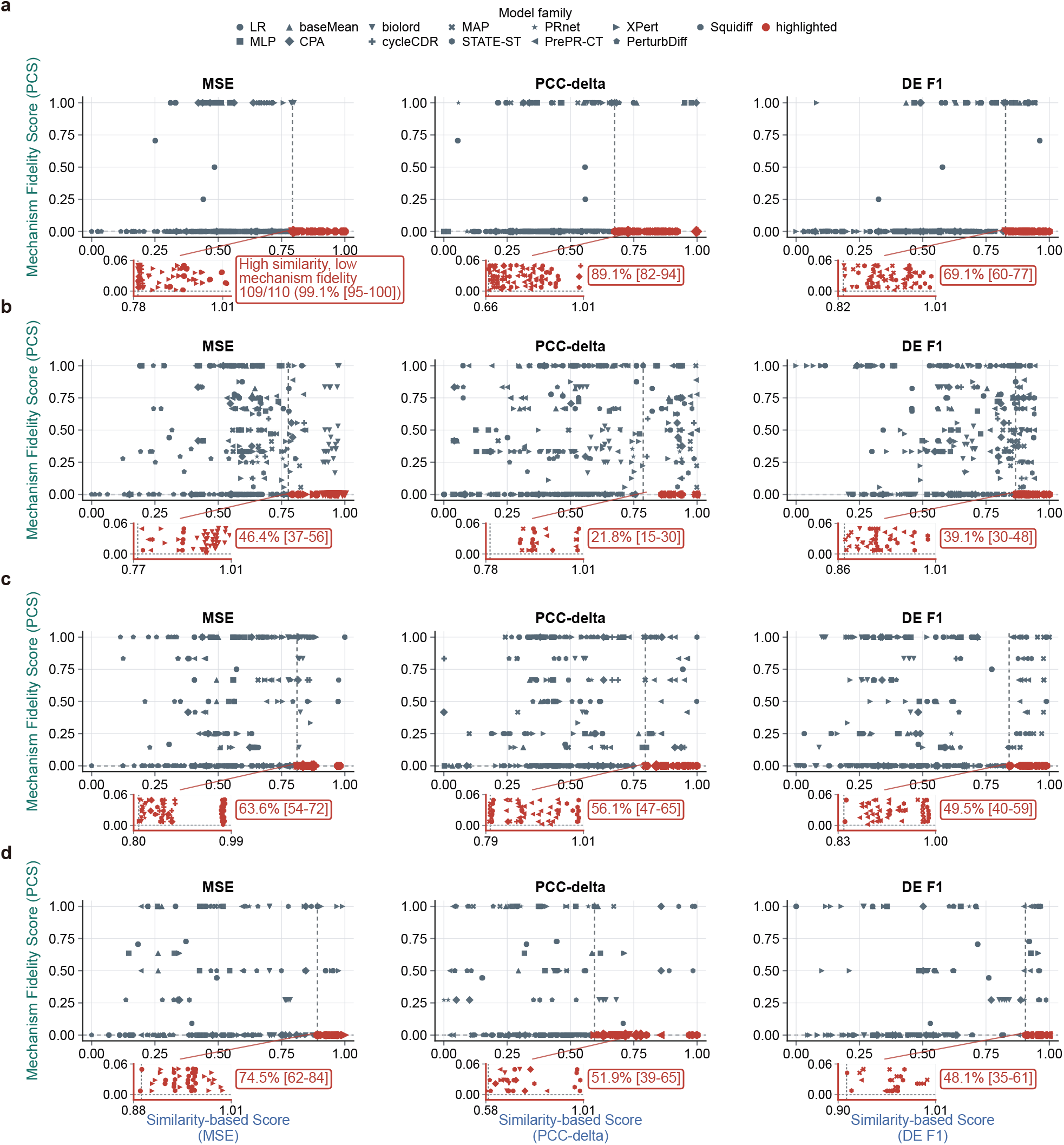
OOD-specific case-level discordance between expression similarity and mechanism fidelity. **a–d**, High-expression-similarity and low-PCS cases corresponding to Fig. 2b across OOD-cell, OOD-drug, OOD-tissue and OOD-drug-cell-pair settings. Expression similarity is shown as the direction-oriented similarity-based score used for visualization, so larger values indicate lower raw error or distance.

**Figure S4.**
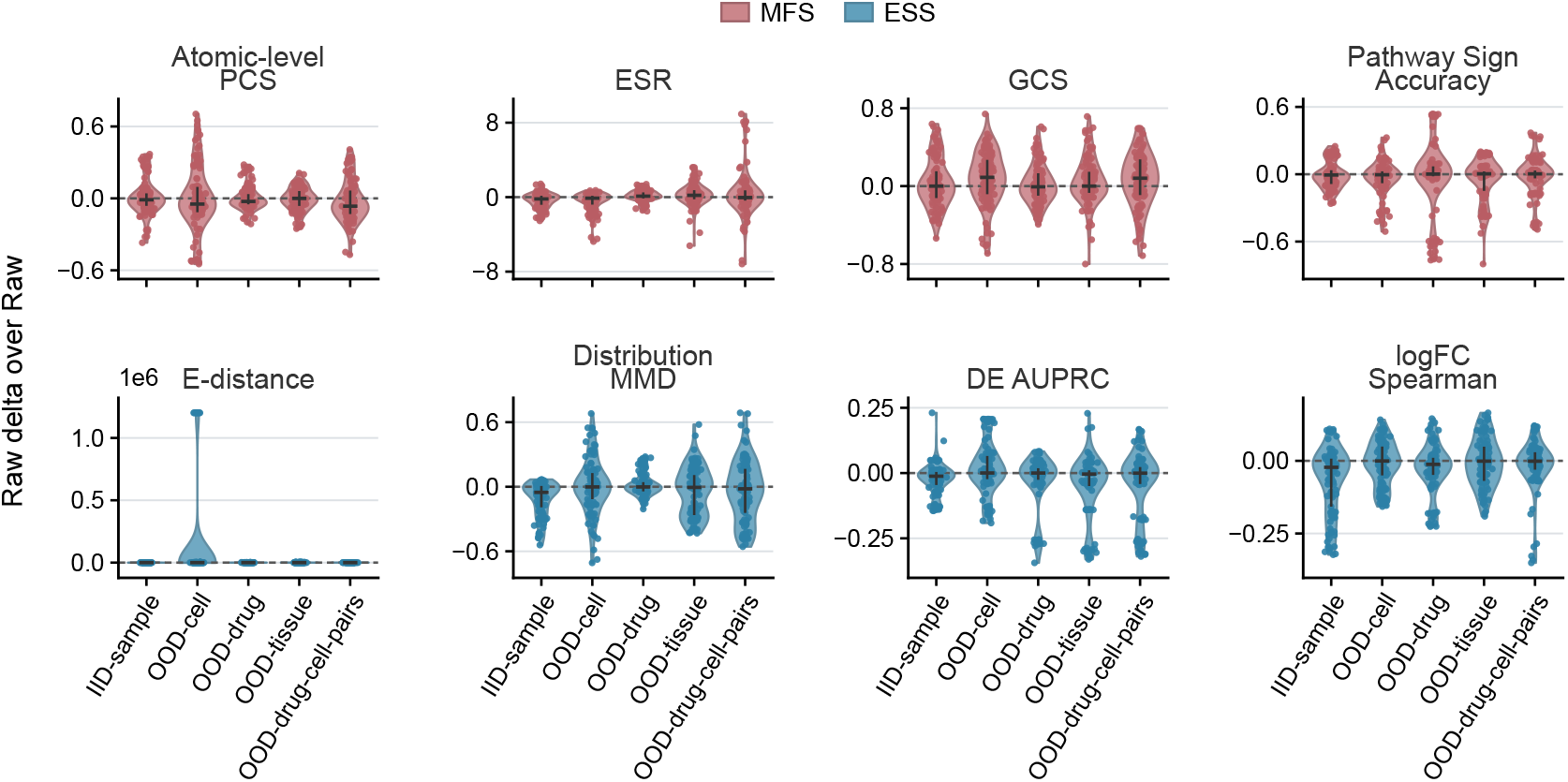
Supplementary raw-delta analysis of representation-augmented cell-line perturbation prediction, complementing Fig. 4d and including Atomic-level PCS, ESR, Pathway Sign Accuracy, E-distance, DE AUPRC and DE F1. Direction-oriented raw-delta distributions for selected mechanism-fidelity and expression-similarity metrics across iid-sample, OOD-cell, OOD-drug, OOD-tissue and OOD-drug-cell-pair settings. Values compare representation-augmented pipelines with matched Raw-input predictors; positive values indicate improved performance relative to Raw input.

**Figure S5.**
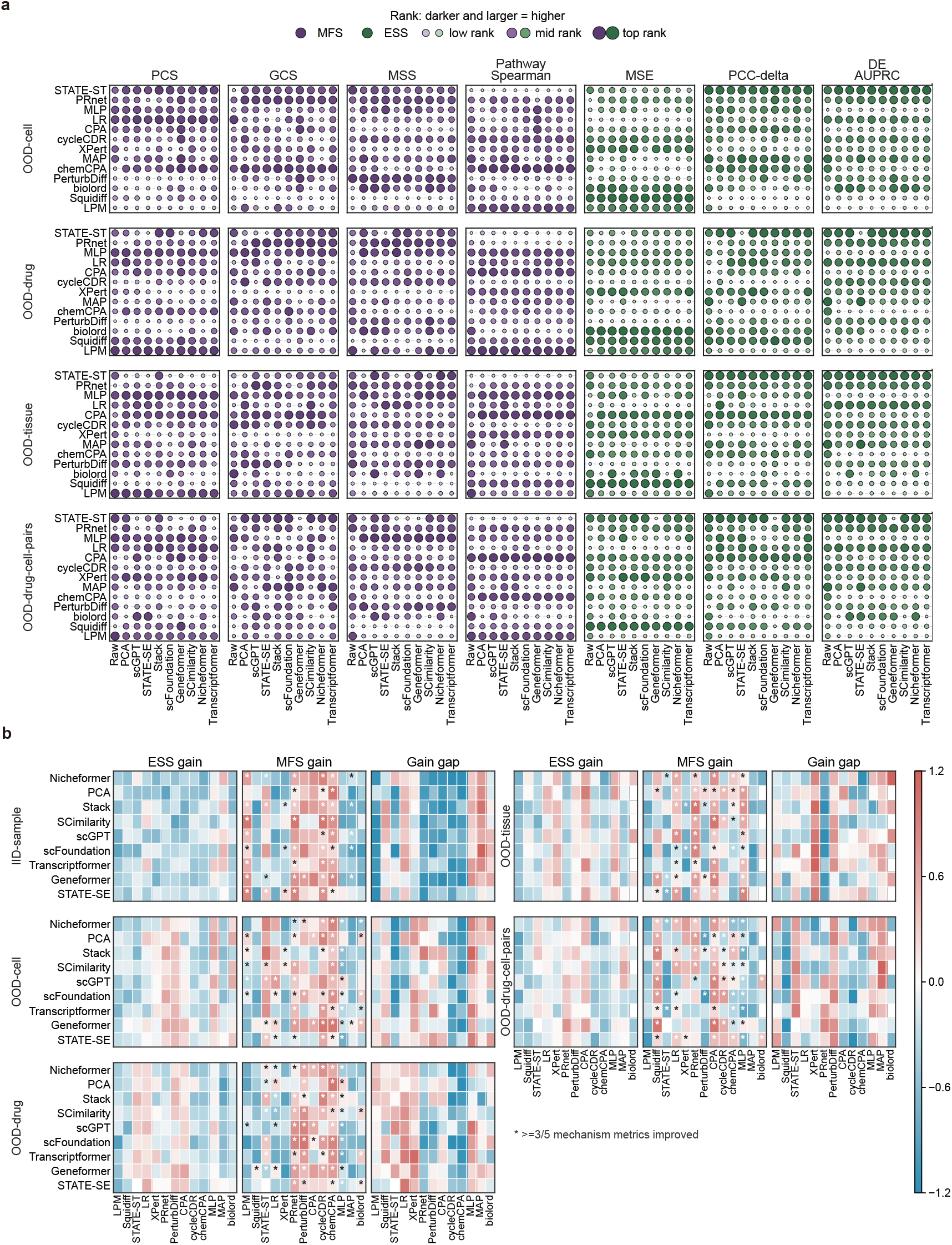
Supplementary analyses of representation-dependent gains in cell-line perturbation prediction. **a**, OOD ranking landscape of scFM embeddings and perturbation predictors across mechanism-fidelity and expression-similarity metrics, complementing the iid-sample ranking analysis in Fig. 4c. **b**, Five-split OOD embedding–predictor gain maps, complementing Fig. 4e and showing expression gain and PCS-prioritized MFS gain in mechanism fidelity relative to Raw input for each scFM or PCA representation paired with each perturbation predictor.

**Figure S6.**
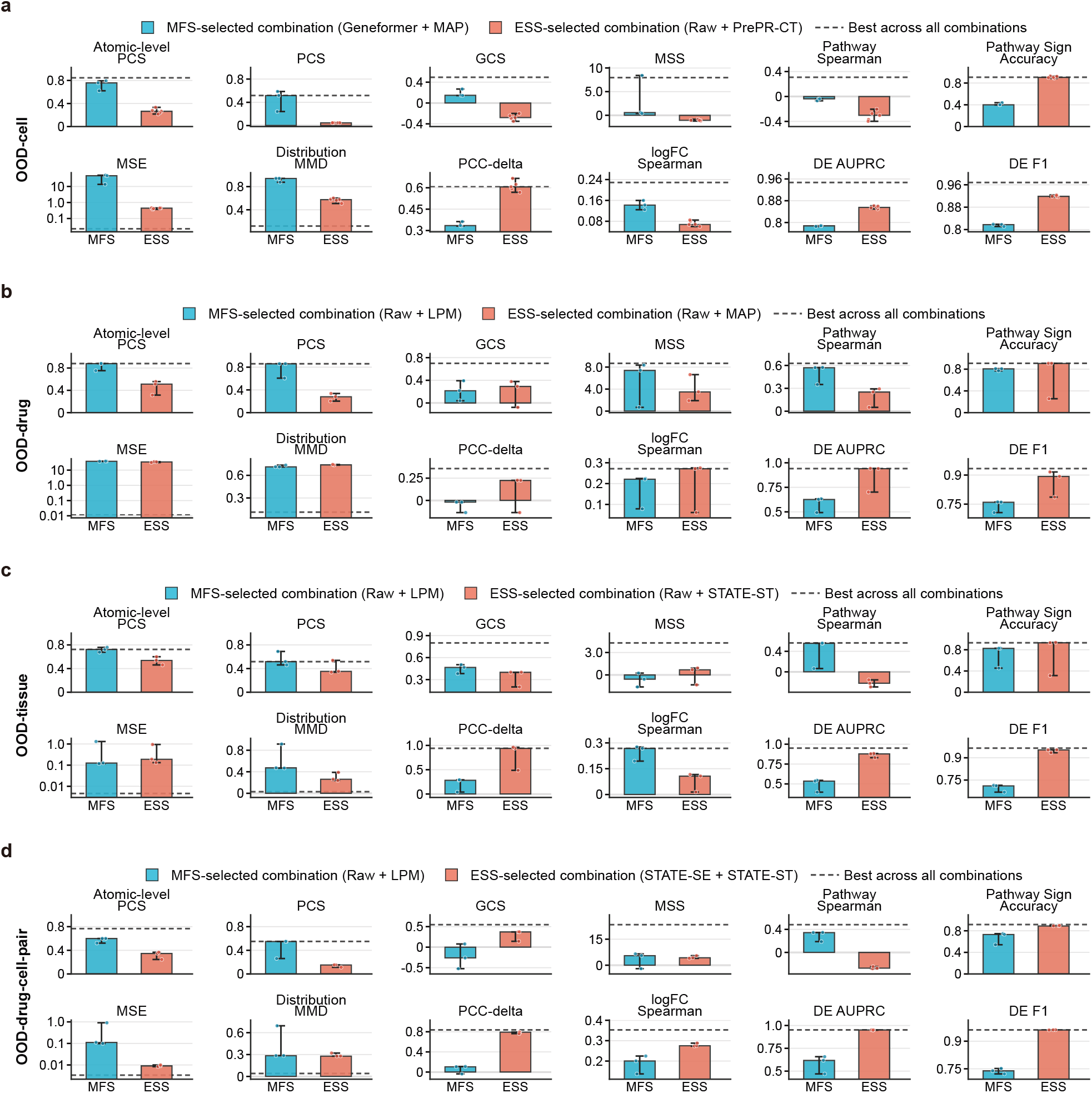
OOD stability of PCS-prioritized MFS- and ESS-selected best combinations across evaluation metrics. **a–d**, Metric-wise performance of the best combinations selected by PCS-prioritized MFS or ESS in OOD-cell, OOD-drug, OOD-tissue and OOD-drug-cell-pair settings, complementing the iid-sample analysis in Fig. 4f. Blue bars indicate MFS-selected combinations and salmon bars indicate ESS-selected combinations. Dashed lines mark the best score achieved by any combination for each metric within the corresponding split.

**Figure S7.**
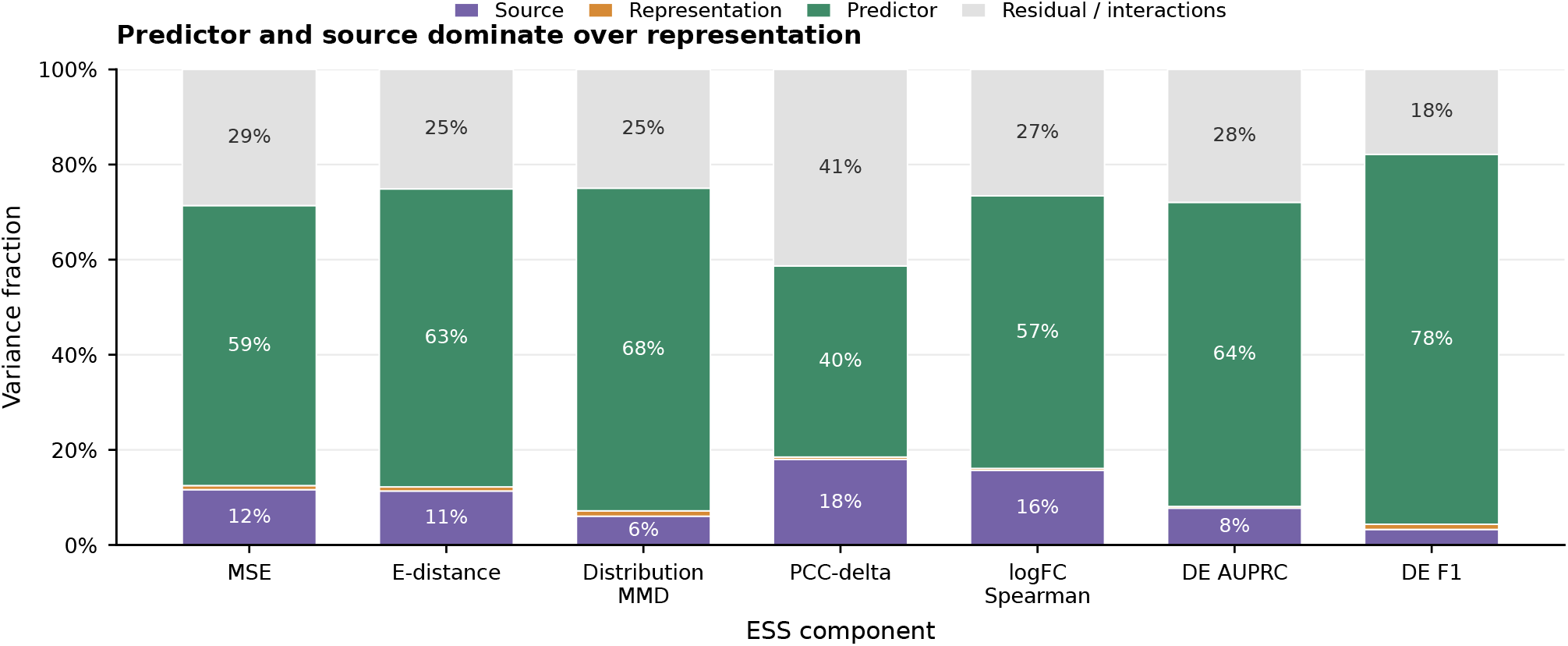
Predictor and source dominate over representation main effects. Variance partitioning of seven expression-similarity score (ESS) metrics using a Type II analysis of variance based on the additive model *Y* ∼ source + representation + predictor. Metric values were percentile-normalized across the complete balanced design comprising five sources, ten representations and thirteen predictors (*n* = 650 observations per metric). Stacked bars show the variance fractions attributed to source, representation, predictor and residual variation. Residual variation includes unmodelled interactions and measurement noise. Predictor and source generally account for larger fractions than the representation main effect, indicating that model architecture and source context have greater influence on performance than the choice of representation alone.

**Figure S8.**
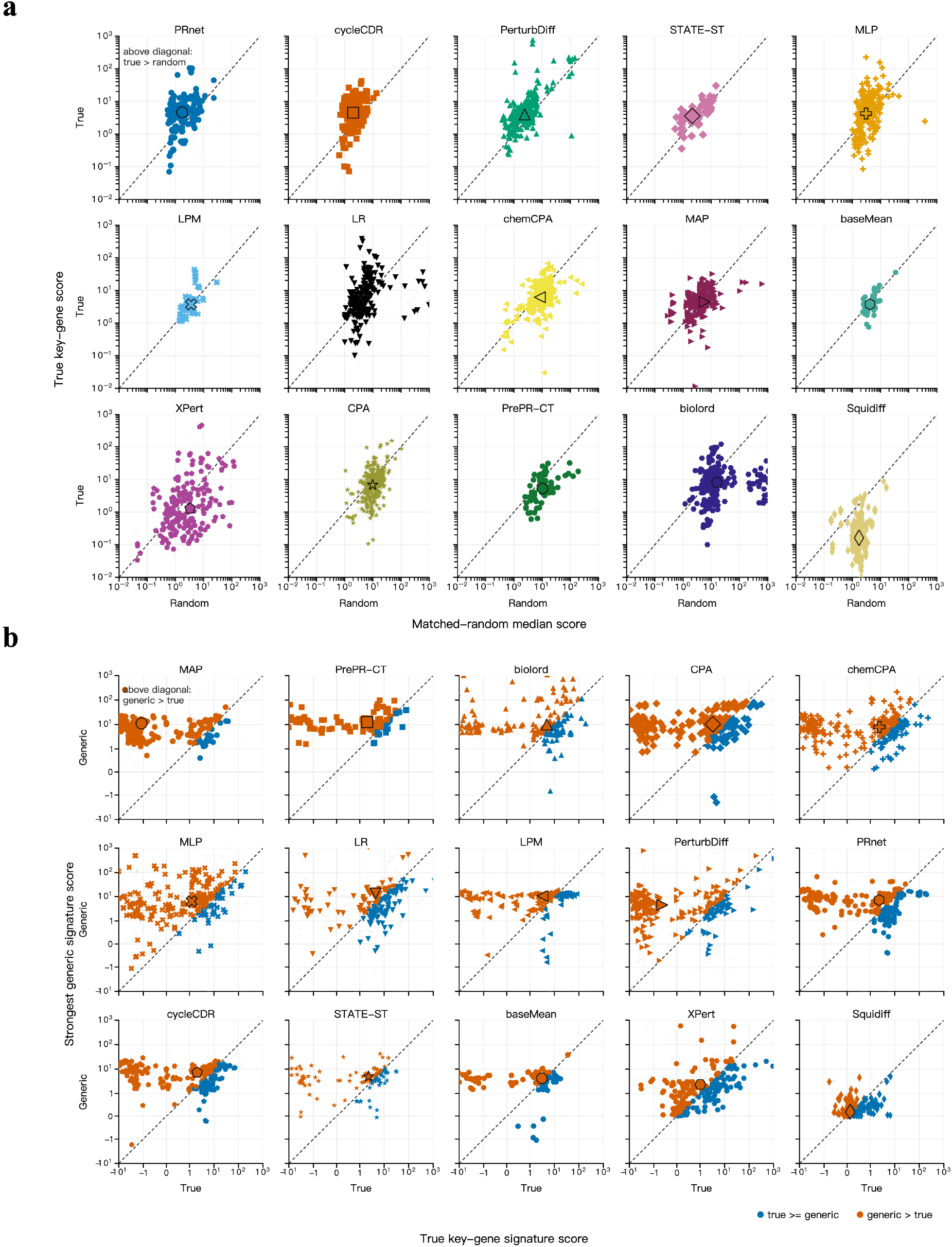
Case-level hard-negative analyses of key-gene specificity and generic-response shortcuts. **a**, Case-level comparison corresponding to Fig. 6b, showing predicted support for true key genes versus expression-matched random genes for each perturbation-prediction model. Points above the diagonal indicate cases in which true key genes received stronger support than matched random genes, whereas points below the diagonal indicate loss of key-gene specificity. **b**, Case-level comparison corresponding to Fig. 6d, showing support for true key-gene signatures versus the strongest generic drug-response signatures. Points above the diagonal indicate cases in which the generic signature exceeded the true case-specific signature, revealing generic-response shortcut behaviour.

**Figure S9.**
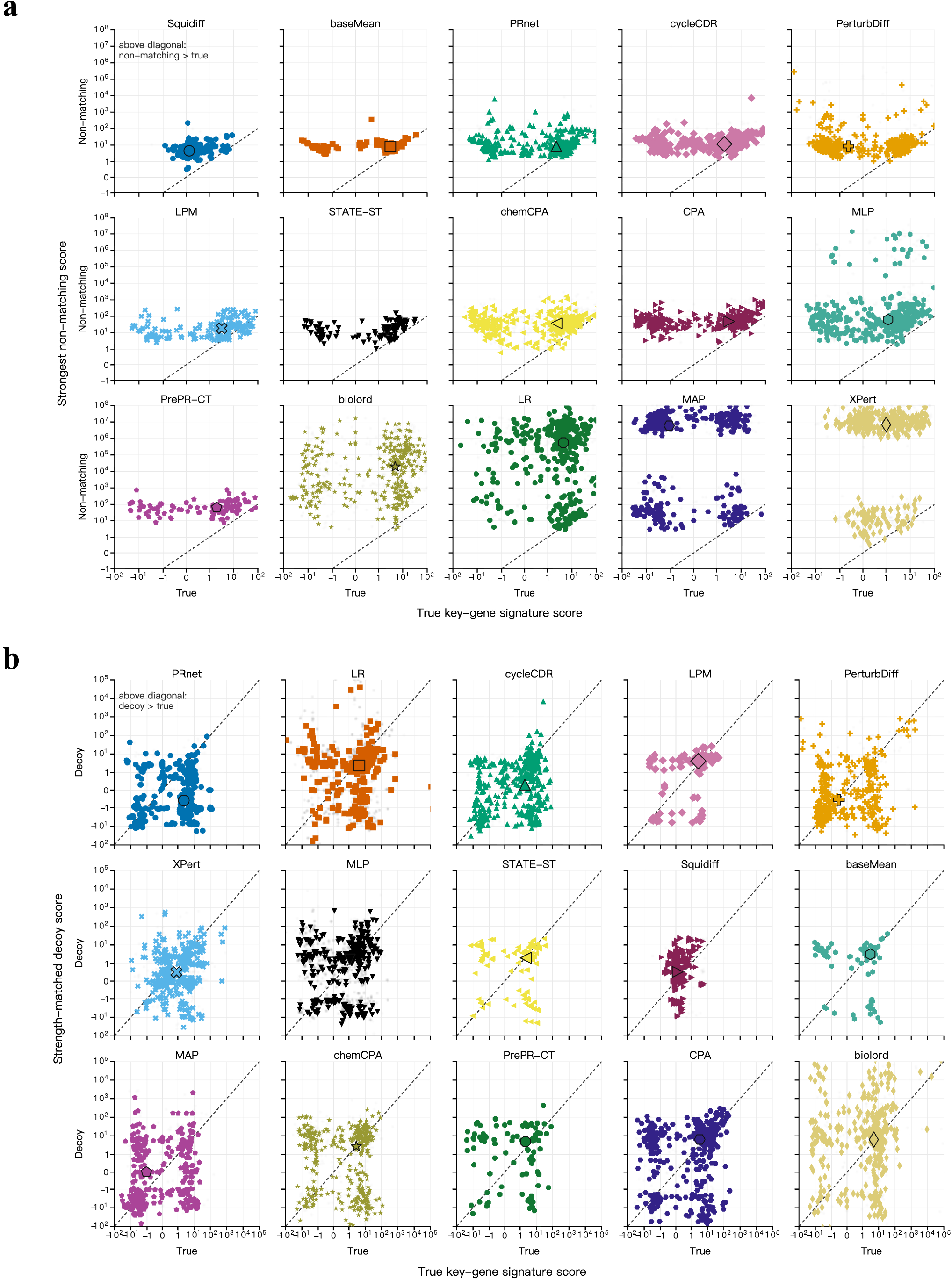
Case-level hard-negative analyses of signature specificity and strength-controlled discrimination. **a**, Case-level comparison corresponding to Fig. 6e, showing support for the true key-gene signature versus the strongest non-matching decoy signature from other perturbation cases. Points above the diagonal indicate cases in which a non-matching signature received stronger support than the true signature, revealing loss of case-specific signature fidelity. **b**, Case-level comparison corresponding to Fig. 6f, showing true key-gene signature scores versus strength-matched negative signature scores for each model. Points below the diagonal indicate cases in which the true perturbation signature scored higher than its strength-matched decoy; points above the diagonal indicate failure to distinguish a drug-specific signature from a response of similar global strength.

## A. Supplementary Methods

This section provides formal definitions and implementation details for the mechanism-fidelity and expression-similarity metrics, composite ranking procedures and hard-negative analyses summarized in the main Methods.

### A.1. Evaluation parameters

Unless otherwise stated, evaluation parameters were fixed before model comparison and applied identically to all models, datasets and data splits. For MFS, bootstrap confidence intervals used *B*_bootstrap_ = 2,000 resamples at a 95% confidence level. In each bootstrap iteration, perturbed cells and matched-control cells were resampled separately with replacement within the corresponding gene–dose unit before recalculating Hedges’ *g*. PCS used these intervals for confidence gating. ESR used *τ*_ESR_ = 0.2 as the minimum absolute observed Hedges’ *g* required for stable predicted-to-observed effect-size ratios. MSS used *R* = 1,000 size-matched random gene sets per test case with random seed 42. These random sets matched the number of target gene–dose units but were not additionally matched by baseline expression. If the null MAD was zero, MSS was set to 0 when the target effect did not exceed the null median and treated as undefined otherwise.

For ESS and pathway-level metrics, log fold changes used a numerical-stability constant *ϵ* = 10^−6^ in both numerator and denominator. Differential-expression recovery used *τ*_DE_ = 0.25 on the absolute log2 fold-change scale. Distribution MMD used an RBF kernel. Unless a bandwidth was specified, *σ* was selected separately for each test case as the median non-zero pairwise Euclidean distance among pooled predicted and observed cell-expression vectors; if all distances were zero, *σ* was set to 1. Pathway metrics used the KEGG 2021 Human gene-set collection in GMT format. The collection contained 320 human pathways represented by gene symbols; genes absent from the evaluated expression matrix were ignored, and no additional pathway-size filter was applied. Pre-ranked GSEA was implemented as a self-contained running-sum procedure using gene-level log fold changes ranked in decreasing order. Enrichment scores were weighted by absolute log fold change and normalized against *B*_GSEA_ = 1,000 size-matched random gene sets to obtain NES values. Pathway Sign Accuracy used *τ*_pathway_ = 1.0 to define observed perturbed pathways. GSEA was used to compute ES and NES values only; no enrichment *P* values were used in the benchmark metrics.

### A.2. Embedding generation details

Frozen scFM and PCA representations differed in how they converted processed expression matrices into cell-level embeddings. These differences included the output layer, pooling rule, expression-tokenization procedure, missing-gene handling and alignment to each model’s gene space (Supplementary Table S1). We therefore generated embeddings with the released implementation and checkpoint-specific preprocessing for each representation model, and used the resulting fixed cell-level vectors as inputs to compatible downstream perturbation-prediction models.

### A.3. Mechanism Fidelity Score

The Mechanism Fidelity Score (MFS) was designed to evaluate whether a predicted perturbation response preserved the annotated drug-response signature. It comprises six complementary metrics. At the gene level, PCS measures response direction, ESR measures effect-size calibration, GCS measures coordinated gene-set structure and MSS measures enrichment of predicted effects in mechanism-relevant genes. At the pathway level, Pathway Spearman measures concordance of enrichment profiles and Pathway Sign Accuracy measures agreement in pathway polarity.

All mechanism metrics followed the same hierarchy from gene–dose units to cases, datasets and the complete benchmark. The atomic unit was a gene–dose pair (*g, d*), where *g* denotes a gene and *d* denotes a dose condition. For source *q* ∈ {real, pred}, let 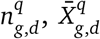 and 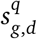 denote the number, mean and sample standard deviation of the cell-level expression values 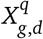 . The corresponding matched-control quantities are 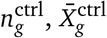 and 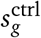. The pooled standard deviation was

**Table S1.** Embedding-generation details for scFM and PCA representations. The table summarizes the model-specific choices used to generate fixed cell-level embeddings before downstream perturbation-prediction benchmarking.

| Representation | Dim. | Layer and pooling | Missing-gene handling | Gene-space alignment |
| --- | --- | --- | --- | --- |
| scGPT | 512 | Last hidden layer; CLS token at position 0; L2 normalization. | Out-of-vocabulary genes were discarded, zero-expression cells were removed and non-zero expression values were binned. | Exact gene-symbol lookup in `vocab.json`; non-zero gene tokens were capped at 1,200 while retaining the CLS token. |
| PCA | 50 | No pretrained layer; PCA fitted on highly variable genes and used to project cells. | Genes outside the highly variable gene subset were not used; the number of PCs was reduced when fewer than 50 highly variable genes were available. | Ground-truth matrices were re-ordered to the matched-control `var_names`; PCA was computed on the shared highly variable gene subset. |
| Geneformer | 1,152 | Second-to-last hidden layer (`emb_layer=-1`); CLS embedding; no L2 normalization. | Genes not mapped to Ensembl identifiers or absent from the token dictionary were discarded; empty cells were removed; sequences were truncated to 4,094 tokens. | Uppercase gene symbols were mapped to Ensembl identifiers, deduplicated and encoded as median-scaled rank-value token sequences. |
| Nicheformer | 512 | Last encoder layer; first three context tokens removed; mean pooling over gene tokens; no L2 normalization. | No explicit out-of-vocabulary filtering was applied; token id was assigned as input-column index plus 30; only the top 1,497 non-zero genes per cell were retained. | Alignment used column position only and therefore depended on the input matrix following the model gene-order convention. |
| scFoundation | Usually 2,048 | Last encoder layer; concatenation of last token, second-to-last token, max-pooled gene tokens and mean-pooled gene tokens; no L2 normalization. | Missing genes were zero-filled in the 19,264-gene canonical space; zero-expression genes did not enter the packed token sequence. | Input genes were reordered to the 19,264-gene scFoundation canonical gene-symbol list. |
| SCimilarity | 128 | Final linear MLP output; direct whole-vector encoder; L2 normalization. | Out-of-vocabulary genes were discarded, missing vocabulary genes were zero-filled and datasets with fewer than 5,000 overlapping genes were rejected. | Exact gene-symbol matching to `gene_order.tsv`; inputs were reordered and zero-filled to the 28,231-gene canonical order. |
| Stack | 1,600 | Final transformer residual stream flattened across the token grid; no pooling and no L2 normalization. | Genes outside the model gene list were discarded, missing positions were zero-filled and inputs with no matched target genes were rejected. | Uppercase exact gene-symbol matching to the checkpoint gene-list pickle; matched genes were placed in the sorted model gene order. |
| STATE-SE | 128 | Final CLS-position output after the decoder stack; L2 normalization. | Genes absent from the protein-embedding vocabulary were masked and discarded; no unknown token was used. | The best-matching gene column was selected automatically, followed by exact string matching to protein-embedding keys. |
| Transcriptformer | 2,048 | Last encoder output; mean pooling over gene tokens; no L2 normalization. | Genes failing symbol-to-Ensembl mapping or vocabulary filtering were discarded; files with no matched genes were skipped; tokenizer fallback used an `unknown` token. | Gene symbols were mapped to Ensembl identifiers with MyGene, Ensembl version suffixes were stripped and identifiers were matched exactly to the model vocabulary. |

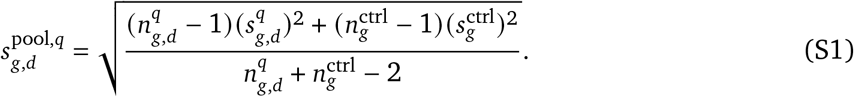

For 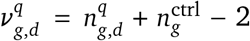 degrees of freedom, the small-sample correction was approximated by *J* (*v*) = 1 − 3/(4*v* − 1). Hedges’ *g* was then defined as

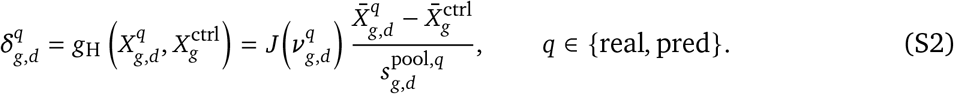

Positive and negative values indicate increased and decreased expression relative to matched control, respectively. These effect sizes were used by all gene-level mechanism metrics below.

#### Pattern Consistency Score (PCS)

PCS quantified whether predicted perturbation effects preserved the observed direction of annotated gene responses.

#### Atomic-level PCS

For each gene–dose unit, uncertainty in 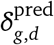 was estimated by cell-level bootstrap resampling with *B*_bootstrap_ = 2,000, yielding a 95% confidence interval:

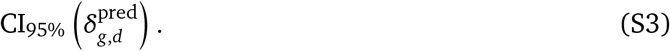

Predictions were considered confident when this interval did not include zero. PCS was then defined as

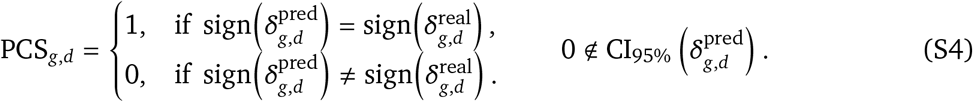

Units with confidence intervals that included zero were treated as inconclusive and excluded from PCS aggregation.

#### Test case-level PCS

Each test case *i* contained a set of annotated gene–dose units:

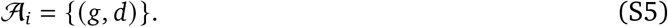

Let the subset of confident units be

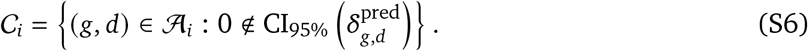

Case-level PCS used a strict consistency rule:

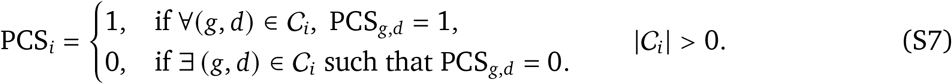

Thus, a case was counted as directionally correct only when all confident units had the correct sign. Cases with |*C*_*i*_| = 0 were unevaluable and excluded from PCS aggregation. We also computed the confident rate:

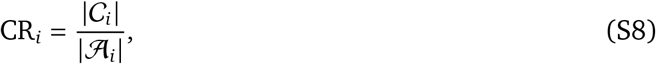

which records the fraction of annotated units that were evaluable.

#### Dataset-level aggregation

Within each dataset, PCS was computed as a weighted mean across valid cases. For dataset *j*, let

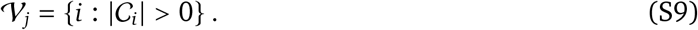

The dataset-level PCS is

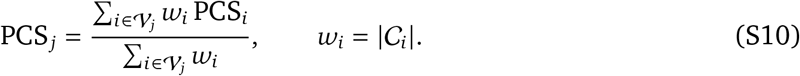

The weight *w*_*i*_ gives greater influence to cases with more confident annotated units. Dataset-level confident rates were summarized as

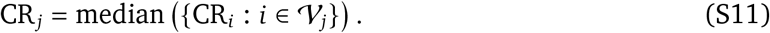

##### Benchmark-wide aggregation

Across datasets J, the overall PCS was defined as

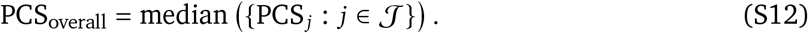

Similarly,

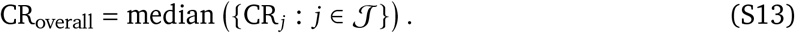

Median aggregation reduces sensitivity to outlier datasets and heterogeneous task difficulty. Confidence gating prevents near-zero, statistically unsupported effects from contributing to directional evaluation.

#### Effect Size Recovery (ESR)

ESR quantified calibration of perturbation-effect magnitude after directional consistency had been established. For each evaluable gene–dose unit, ESR was defined as the ratio of predicted to observed effect size:

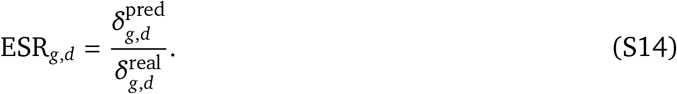

ESR was computed only for confident and directionally correct units, PCS_*g,d*_ = 1, with sufficiently large observed effects, 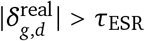, where *τ*_ESR_ = 0.2.

For cases involving multiple genes or doses, each gene–dose pair was treated as an evaluable unit. Let

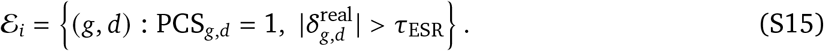

Case-level ESR was defined as

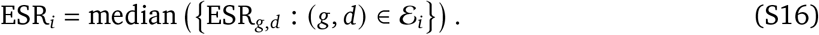

The median was used to limit the influence of unstable ratios.

For each dataset *j*, ESR was aggregated across valid cases:

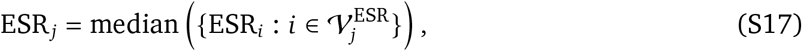

and variability was quantified as

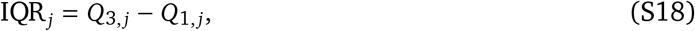

where 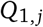 and 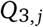 are the 25th and 75th percentiles of 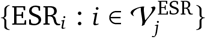. Aggregating at the case level prevents cases with more annotated units from dominating the dataset summary.

To jointly summarize magnitude accuracy and stability across the benchmark, we used an overall ESR score:

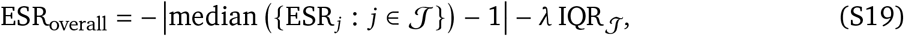

where IQR_*J*_ is the interquartile range of dataset-level ESR values. The dispersion penalty λ was set to 0.5 by default. Higher values indicate smaller deviation from the ideal ratio of 1 and lower between-dataset variability.

#### Gene-set Coherence Score (GCS)

GCS evaluated whether a model preserved coordinated responses within an annotated gene set. For each test case *i* 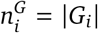 and 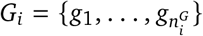. We formed predicted and observed effect-size vectors under condition *d*_*i*_:

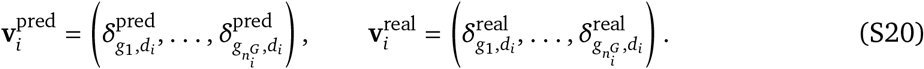

GCS was defined as the Spearman rank correlation between these vectors:

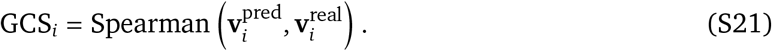

GCS was computed only for gene sets with 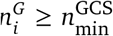. Case-level GCS values were summarized by the median within each dataset *j*:

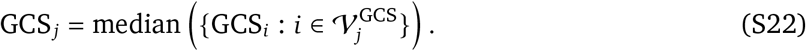

Overall GCS was then defined as the median across dataset-level scores:

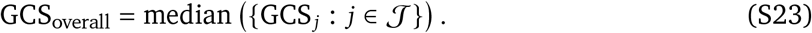

Variability was quantified by the interquartile range of dataset-level scores. GCS therefore measures preservation of relative response ordering within a gene set, rather than only aggregate effect magnitude.

For mean-based summaries and statistical analyses, correlation coefficients were transformed using Fisher’s *z*:

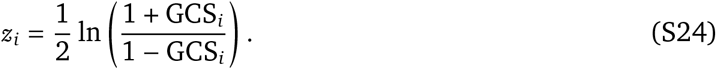

For summary statistics, we computed

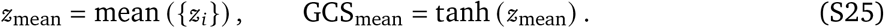

The median GCS was used as the primary summary. The Fisher-transformed mean was reported as a supplementary statistic.

#### Mechanism Specificity Score (MSS)

MSS evaluated whether predicted effects were enriched in mechanism-relevant genes rather than distributed across background genes. For each test case *i* with target gene set *G*_*i*_ and dose condition *d*_*i*_, the aggregate target effect was

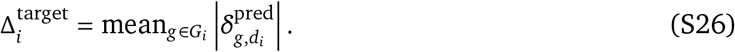

To construct a background distribution, we sampled *R* = 1,000 size-matched random gene sets 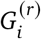 and computed

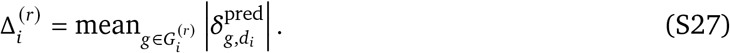

MSS was defined as a robust standardized contrast:

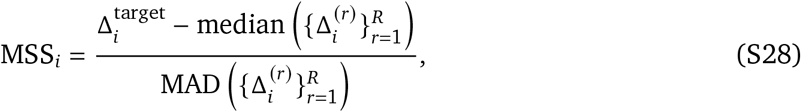

with zero-MAD null distributions handled as described in the Evaluation parameters section. MAD denotes the median absolute deviation:

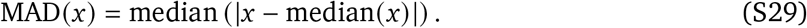

Across test cases, MSS was summarized by the median:

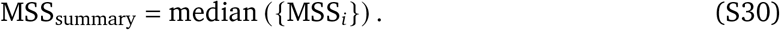

Positive MSS values indicate stronger predicted effects in the annotated target genes than in matched random gene sets.

#### Pathway Spearman

Pathway Spearman quantified concordance between predicted and observed pathway-response profiles. For each perturbation *p* and gene *g*, log fold changes were computed from mean expression in the predicted or observed perturbed condition relative to control:

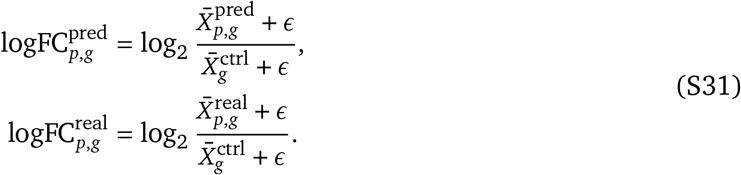

where 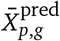 and 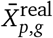 are the mean expression levels of gene *g* under perturbation *p* in the predicted and observed data. 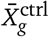 is the corresponding control mean.

Predicted and observed logFC vectors were analyzed independently by pre-ranked gene-set enrichment analysis against the KEGG 2021 Human pathway collection, denoted *P*= *P*_1_, …, *P*_*K*_ . For pathway *P*_*k*_, genes were ranked by decreasing logFC and scored with an absolute-logFC-weighted running-sum enrichment statistic. Each enrichment score was normalized against *B*_GSEA_ = 1,000 size-matched random gene sets to obtain a normalized enrichment score (NES). Pathway Spearman was then defined as the Spearman correlation between predicted and observed NES values over pathways with finite scores in both conditions:

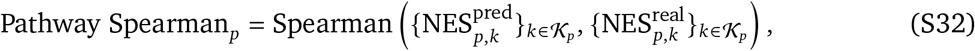

where 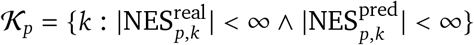.

#### Pathway Sign Accuracy

Using the same NES values, a pathway was considered perturbed if its observed NES magnitude exceeded *τ*_*pathway*_ = 1.0. Let 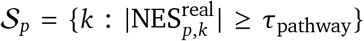 denote this pathway set. Pathway Sign Accuracy was defined as the fraction of these pathways for which predicted and observed enrichment polarity agreed:

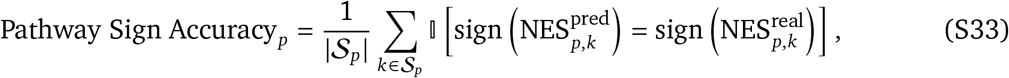

where sign (·) returns +1 for positive values and −1 for negative values, and I[·] is the indicator function. If *S*_*p*_ = ∅, the metric was reported as undefined for that perturbation. This metric isolates pathway polarity from the rank-order concordance captured by Pathway Spearman.

### A.4. Expression-similarity metrics

Conventional transcriptomic similarity was evaluated using seven metrics. For *q* ∈ {real, pred }, the mean expression of gene *g* under condition *d*_*i*_ was

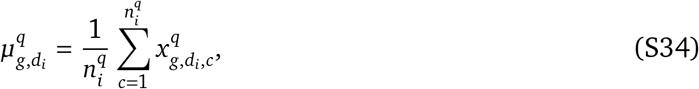

where 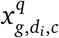 is the normalized expression value in cell *c*, and 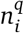 is the number of cells. Mean perturbation effects were computed relative to the matched-control mean:

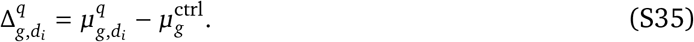

Log fold changes were defined separately as

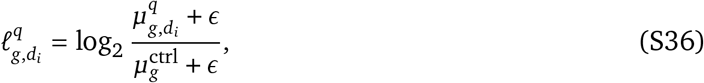

where *ϵ* = 10^−6^ is a numerical-stability constant.

#### Mean Squared Error (MSE)

MSE quantified the average squared discrepancy between predicted and observed perturbation effects:

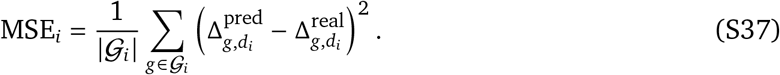

#### E-distance

E-distance compared the full predicted and observed single-cell populations. Let 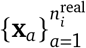 and 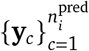 denote their cell-expression vectors over G_i_ . The empirical E-distance was

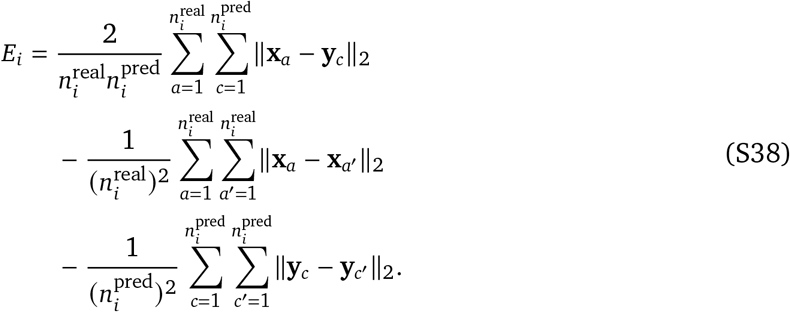

#### PCC-delta

PCC-delta measured Pearson correlation between the predicted and observed mean perturbation-effect vectors:

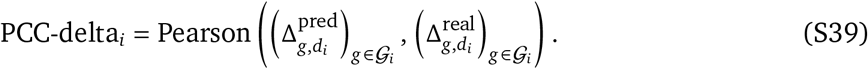

#### DE AUPRC

DE AUPRC evaluated differential-expression recovery as a ranking problem. Observed DE labels and predicted scores were defined as

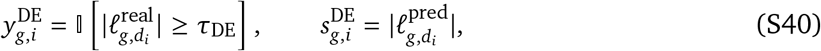

and the case-level score was

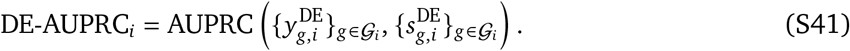

#### DE F1

DE F1 treated both observed and predicted DE status as binary calls at the same threshold. With 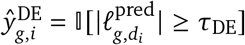, it was defined as

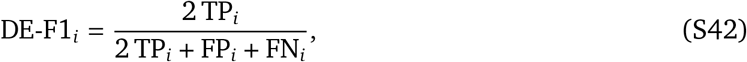

where TP_*i*_, FP_*i*_ and FN_*i*_ were calculated from 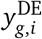 and 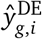 over *G*_*i*_.

#### logFC Spearman

logFC Spearman measured whether the rank ordering of gene-level perturbation effects was preserved:

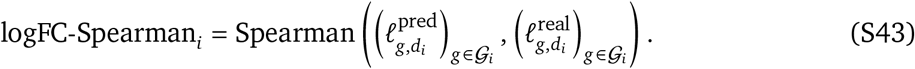

#### Distribution MMD

Distribution MMD compared predicted and observed cell populations using the RBF kernel

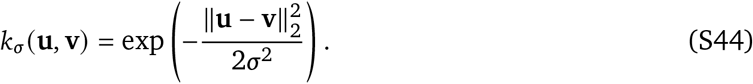

The unbiased empirical squared maximum mean discrepancy was

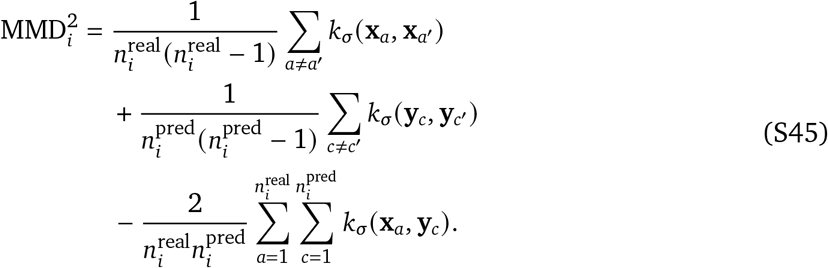

Lower MSE, E-distance and MMD values indicate better agreement.

Unless otherwise specified, each metric was first computed at the test-case level. For a generic metric *M*_*i*_, let 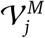 denote its valid test cases within dataset *j*. The median was the primary dataset-level summary and the value used to construct ESS. Arithmetic means were retained only as supplementary descriptive statistics:

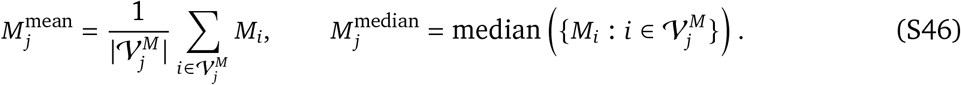

Overall benchmark results were summarized by the median across datasets *J* MMD used the median pairwise-distance heuristic to select *σ*, computed from pooled predicted and observed cell-expression vectors. Metrics with undefined correlations, empty positive classes or zero denominators were excluded from the corresponding aggregation. The absolute log2 fold-change threshold *τ*_DE_ = 0.25 and numerical-stability constant *ϵ* = 10^−6^ were fixed across model comparisons.

#### Direction-oriented similarity-based scores for case-level visualization

For case-level visual analyses comparing expression similarity with mechanism fidelity, including Fig. 2b and Supplementary Fig. S2c, expression metrics were converted to direction-oriented similarity-based scores within each data split and each metric. This transformation was used for visualization and discordance calling, and was separate from the robust *z*-score aggregation used to construct ESS for model ranking. For MSE and E-distance, each valid raw value *x* was first log-transformed:

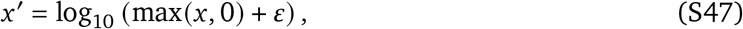

With

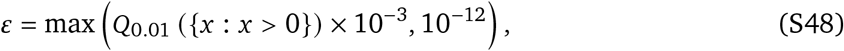

where *Q*_0.01_ is the first percentile of positive valid values for that split and metric. The transformed values were then min–max normalized and reversed:

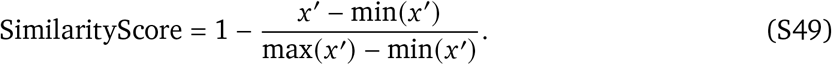

Distribution MMD was not log-transformed and was instead reversed directly after min–max normalization:

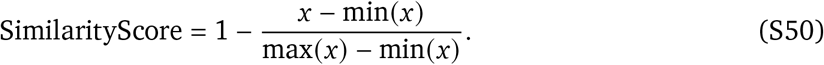

Thus, lower raw MSE, E-distance and Distribution MMD values corresponded to larger similarity-based scores. If all valid values for a metric were identical within a split, all observations were assigned a similarity-based score of 0.5. In the iid-sample validation set, Spearman correlations between raw values and similarity-based scores were − 1.0 for MSE, E-distance and Distribution MMD. High-expression/low-mechanism discordance was defined using the top quintile of these direction-oriented similarity-based scores and the bottom quintile of PCS, equivalently SimilarityScore ≥ *Q*_0.80_ and PCS ≤ *Q*_0.20_ within the corresponding split and metric.

### A.5. Methods ranking

For any dataset–model pair *u* = (*j, m*) and metric *S*, values were first oriented so that larger values indicated better performance:

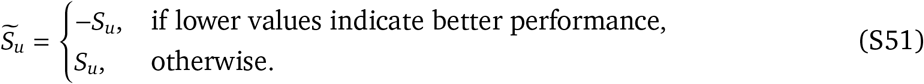

When metrics with different scales were combined, we used a robust *z*-score computed over the valid dataset–model pairs *U*_*S*_:

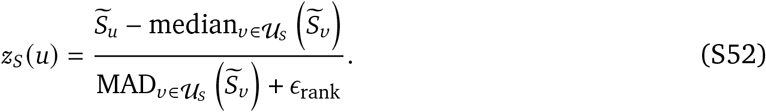

where *ϵ*_rank_ is a small stabilizer used only for robust rank standardization. Median aggregation was used throughout the composite scores to reduce sensitivity to individual metrics with extreme values or nested *K*-dependent definitions.

#### Mechanism-fidelity ranking

Mechanism-fidelity ranking used a PCS-prioritized MFS rule. PCS was used as the primary ordering variable because it directly measures whether the predicted response preserves the annotated direction of key-gene effects. Auxiliary mechanism information was used only to break ties or near-ties in PCS. For each dataset, models were first sorted by PCS in descending order and assigned to PCS tiers using a tolerance *δ*_PCS_ = 0.02. The top remaining model defined the tier anchor, and all remaining models *m* satisfying

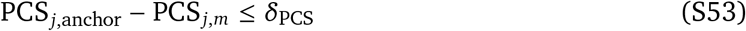

were assigned to the same tier. The procedure was repeated until all models in dataset *j* had been assigned to a tier.

Within each PCS tier, models were ranked by an auxiliary mechanism score computed from gene-set and pathway-level mechanism metrics:

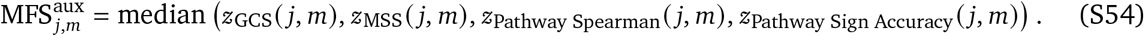

The final MFS ranking was therefore defined by the ordered keys

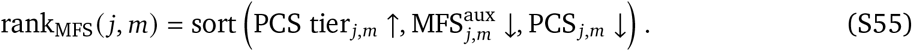

This rule allows GCS, MSS and pathway-level metrics to refine ranking among models with similar directional key-gene consistency, while preventing secondary metrics from overriding a substantial PCS difference. ESR was not included in the primary MFS ranking because it is a ratio-based calibration metric computed only after confident directional consistency. We therefore reported ESR separately and used it for sensitivity analyses of effect-size calibration. PCS, ESR, GCS, MSS, Pathway Spearman and Pathway Sign Accuracy were also reported separately.

#### Expression-similarity ranking

Expression-similarity ranking used an ESS that combined seven conventional transcriptomic similarity metrics. For each expression-similarity metric, the dataset– model score was the median across valid test cases:

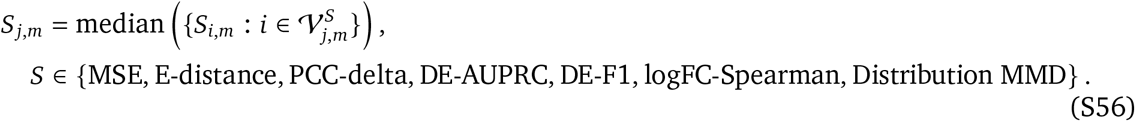

Cases for which a metric was undefined were excluded only from the aggregation of that metric. This procedure gave each valid test case equal weight, irrespective of its cell or gene count.

Error and distance metrics were sign-oriented before standardization so that larger *z*-scores consistently indicated better performance. For each dataset–model pair (*j, m*), the seven metrics were grouped into three component scores:

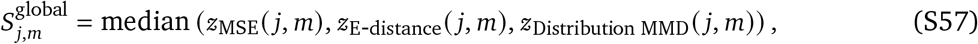

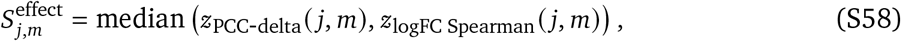

and

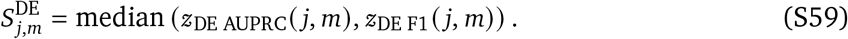

The final ESS was defined as

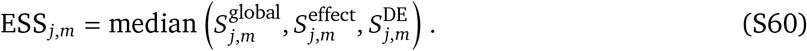

Expression-similarity ranks were assigned by sorting ESS in descending order within each dataset. Pathway Spearman and Pathway Sign Accuracy were excluded from ESS and used only for mechanism-fidelity ranking.

#### Drug-retrieval ranking

Drug-retrieval ranking summarized downstream retrieval performance while emphasizing early enrichment of the true drug among candidate compounds. All drug-retrieval metrics were oriented such that larger values indicated better performance. We first grouped nested *K*-dependent metrics to avoid over-weighting repeated Hit@K or NDCG@K measurements. Early hit enrichment was defined as

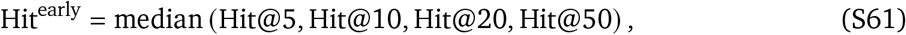

and broad hit recovery was defined as

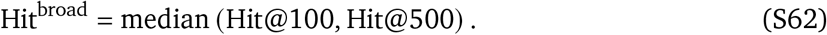

Analogously, early and broad ranking quality were

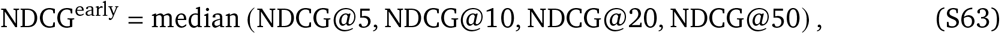

and

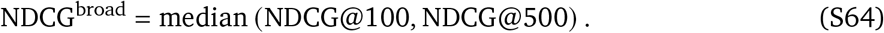

Mean reciprocal rank (MRR) was retained as a separate first-hit component because it is sensitive to the rank of the earliest correct retrieval.

The primary drug-retrieval composite score was the Early Drug Retrieval Score:

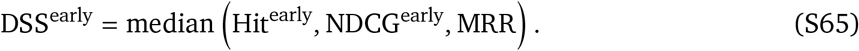

Representations were ranked for downstream drug retrieval by descending DSS^early^. As a sensitivity analysis, we also computed a full-range score,

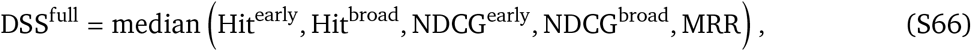

which incorporates broader candidate lists while retaining early retrieval and first-hit information.

### A.6. Source-aware analyses in the multi-source benchmark

The source-aware analysis was designed to test whether mechanism fidelity, expression reconstruction and model selection were stable across biological source contexts. We analysed five splits: IID-sample, OOD-cell-line, OOD-primary-culture, OOD-organoid and OOD-patient-sample. Unless otherwise specified, each pipeline–split value was first summarized as the median across repeated experiments before downstream ranking, gain analysis or variance partitioning.

#### Source-level mechanism-fidelity profiles

For the source-level distribution analysis, we summarized six mechanism-fidelity components separately: Atomic-level PCS, PCS, GCS, MSS, Pathway Spearman and Pathway Sign Accuracy. Pipelines were grouped as Raw/PCA baseline pipelines or scFM-enhanced pipelines. The Raw/PCA group contained 28 pipelines per split, whereas the scFM-enhanced group contained 104 pipelines per split, corresponding to eight frozen scFM representations combined with the common downstream predictors. Each plotted point represented one pipeline median across repeated experiments. Box centres and hinges denoted the median and interquartile range, and whiskers denoted the 5th and 95th percentiles. These plots used the original metric values rather than a composite MFS or rank percentile; therefore, comparisons were interpreted within each metric rather than across metrics with different numerical scales.

#### Cross-split ranking concordance

To assess whether source context reordered pipelines, we compared source-specific rankings for the same matched pipelines. Mechanism-fidelity rankings used the PCS-prioritized MFS rule described above, with *δ*_PCS_ = 0.02. Within a PCS tier, auxiliary ranking used the median robust *z*-score of GCS, MSS, Pathway Spearman and Pathway Sign Accuracy. ESR was not included in this primary source-aware MFS ranking. Expression-similarity rankings used ESS, with MSE, E-distance and Distribution MMD sign-oriented before robust *z*-score standardization.

For each pair of source-aware splits *s* and *t*, ranking concordance was computed as the Spearman correlation between the matched pipeline ranks:

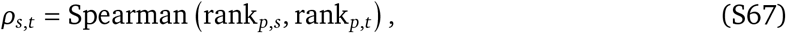

where *p* indexes pipelines present in both splits. The mean concordance of split *s* with the remaining splits was

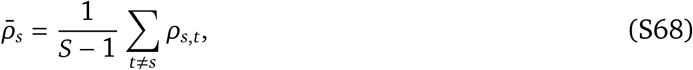

where *S* = 5. In the MFS concordance heatmap, cells with ρ < 0.15 were marked to highlight near-absent rank agreement.

#### Matched scFM gain analysis

To isolate the contribution of frozen scFM representations from the downstream predictor, each scFM-enhanced pipeline was paired with the Raw-input pipeline using the same predictor and source-aware split. PCA was not included in this matched scFM–Raw comparison. With eight scFM representations and 13 common predictors, this gave 104 matched comparisons per split and 520 comparisons across the five splits. For an MFS component *m*, the scFM gain was defined as

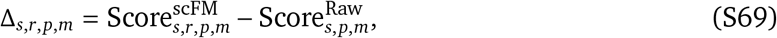

where *s, r, p* and *m* denote source split, scFM representation, predictor and metric, respectively. Because all MFS components are higher-is-better, Δ > 0 indicated improvement.

For ESS metrics, gains were oriented so that positive values always indicated improvement. For lower-is-better metrics, including MSE, E-distance and Distribution MMD, gain was computed as Raw minus scFM. For higher-is-better metrics, including PCC-delta, logFC Spearman, DE AUPRC and DE F1, gain was computed as scFM minus Raw. The positive-gain fraction for each source–metric combination was

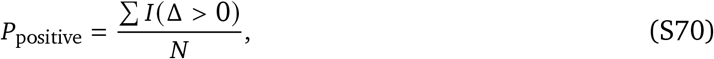

with *N* = 104 matched comparisons per heatmap cell. To assess whether scFM gains were coordinated across evaluation dimensions, we pooled the 520 matched comparisons and computed Spearman correlations between gain vectors for metric pairs:

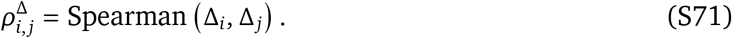

These analyses were used as descriptive summaries of improvement frequency and cross-metric gain concordance, rather than as tests of statistical significance or causal effects.

#### Variance partitioning by source, representation and predictor

We used Type II analysis of variance to estimate the main-effect contribution of source, input representation and downstream predictor to each evaluation metric. To keep the design balanced, we analysed five source-aware splits, ten representations (Raw, PCA and eight scFM embeddings) and 13 predictors that were present for all representations and sources, yielding 650 observations per metric. Each observation was the repeated-experiment median for one source–representation–predictor pipeline.

Because metric scales differed, each metric *m* was transformed independently to a pooled percentile rank across the complete balanced design:

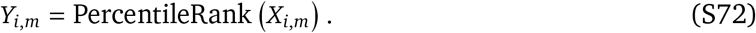

For lower-is-better ESS metrics, values were reversed before percentile transformation so that higher percentiles consistently indicated better performance. Percentiles were computed across the pooled 650 observations rather than within each source, because within-source normalization would remove the source main effect by construction.

For each MFS or ESS component, we fitted the additive fixed-effect model

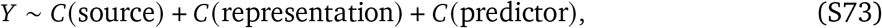

and used Type II sums of squares. The variance fraction attributed to factor *k* was

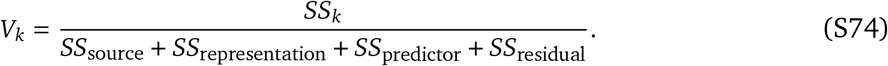

The residual term contains unmodelled interactions, pipeline-specific variation, repeated-experiment variability and other unexplained variation; it was therefore not interpreted as a pure interaction effect. The variance-partitioning analysis was interpreted as an association analysis rather than as evidence of causality.

#### Source-aware model-selection strategies

We compared four model-selection strategies across the same five source-aware splits and matched pipelines. First, ESS-selected pipelines were chosen by maximizing the mean ESS across sources:

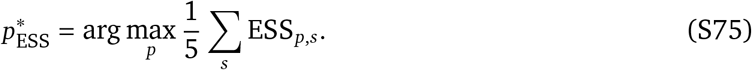

Second, pooled MFS selection summarized each MFS component for a pipeline by its median across the five sources and then applied the PCS-prioritized MFS ranking rule. Third, split-balanced MFS selection computed the MFS rank percentile within each source and then averaged these source-specific percentiles:

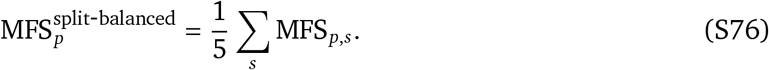

Fourth, worst-split-aware MFS selection used the lowest source-specific MFS percentile for each pipeline:

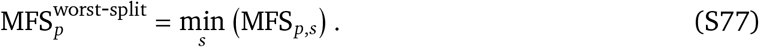

The first strategy emphasizes expression reconstruction, the second emphasizes pooled mechanism fidelity, and the latter two give equal or conservative weight to source-specific mechanism fidelity.

### A.7. Hard negative analysis

Hard negative analysis was designed to identify shortcut solutions that can yield apparently plausible perturbation predictions without preserving the case-specific drug-response signature. Each evaluable unit was a test case *i*, defined by study, cell context, drug, dose and time, and each prediction was indexed by model *m*. For case-level analyses, we used the predicted and observed perturbation-effect vectors

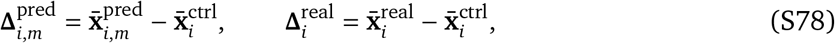

where bars denote pseudobulk mean expression vectors over the aligned gene space. For gene-level mechanism tests, we used the predicted Hedges’ *g*, 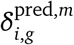, computed between 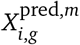 and 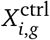 as defined above. The literature-curated mechanism signature for case *i* was denoted *G*_*i*_ = {(*g, s*_*i,g*_)}, where *s*_*i,g*_ = +1 for upregulated genes and *s*_*i,g*_ = −1 for downregulated genes. The corresponding key-gene identity set was *K*_*i*_ = {*g* : (*g, s*_*i,g*_) ∈ *G*_*i*_ }. Genes annotated as non-significant were excluded from direction-based hard negatives.

For any signed gene signature *Q* = {(*g, s*_*g*_)}, we defined a signed mechanism support score

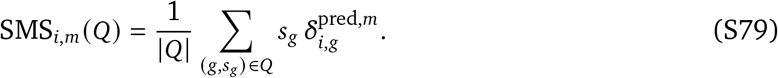

Positive values indicate that the model prediction supports the direction of the signature, whereas negative values indicate support for the opposite direction. To reduce biases caused by signature size, baseline expression and gene-level detectability, signed scores were standardized against expression-matched random signatures:

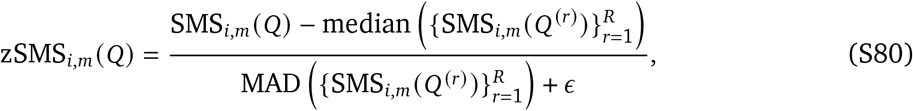

where *Q* ^(*r*)^ has the same size and direction composition as *Q*, and its genes are matched to *Q* by baseline mean expression, detection rate, expression variance and highly variable gene status in the control cells. Unless otherwise specified, *S*_*i,m*_ (*Q*) denotes this standardized score.

#### No-change collapse

We first tested whether the predicted perturbation response collapsed towards the unperturbed state. The no-change ratio was defined as

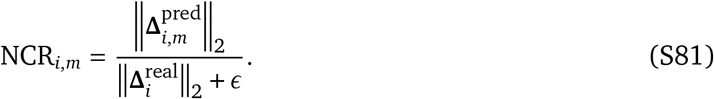

A case was labelled as no-change collapse when NCR_*i,m*_ *<* 0.25, indicating that the predicted response norm was less than one quarter of the observed perturbation norm. Sensitivity analyses used nearby thresholds.

#### Mean-response memorization

We next tested whether predictions were closer to a training-set average perturbation response than to the corresponding observed response. The training-set mean response was estimated as a robust pseudobulk median,

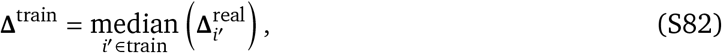

computed separately within each data split. Cosine similarity was used to compare the predicted response with this generic training response and with the true case-specific response:

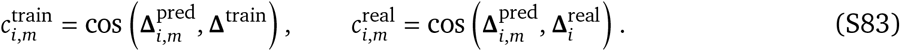

Here, cosine similarity was computed as

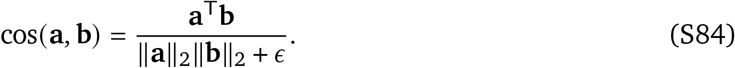

Mean-response memorization was recorded when 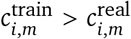.

#### Key-gene specificity against matched random genes

To test whether predicted effects were concentrated on mechanism-relevant genes rather than expression-matched background genes, we compared the mean absolute predicted effect over true key genes with a null distribution from matched random gene sets. For the unsigned key-gene score,

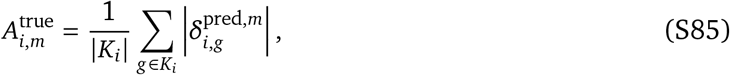

where *K*_*i*_ is the key-gene identity set. For each case, *R* random gene sets 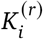 were sampled with the same set size and matched control-expression properties. The corresponding null scores were

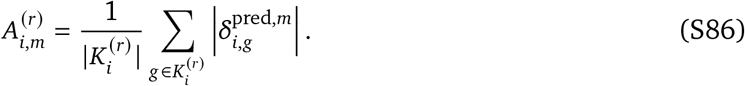

The hard-negative specificity score was the same robust contrast used for MSS:

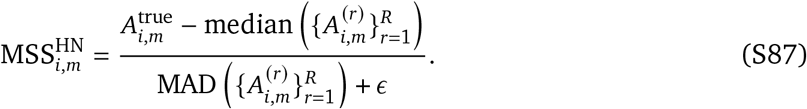

We also recorded a random-rejection indicator,

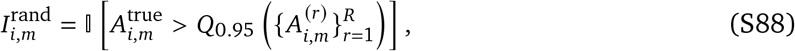

which equals one when the true key-gene score exceeds the 95th percentile of the matched-random null distribution.

#### Direction-flipped decoys

Directional consistency was assessed by comparing the true signed key-gene signature with a decoy in which gene identities were preserved but all directions were reversed:

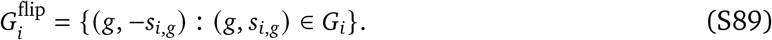

Predicted gene directions were confidence-gated using bootstrap confidence intervals for 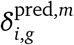. Let

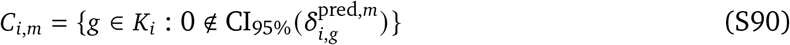

be the confident key-gene set. True and flipped directional consistency were then

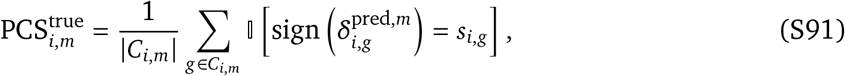

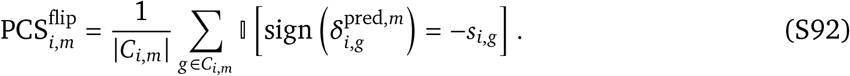

Cases with |*C*_*i,m*_| = 0 were treated as not evaluable for this test. The direction rejection score was

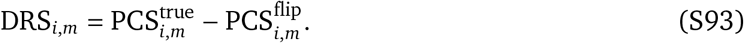

A direction failure was recorded when DRS_*i,m*_ ≤ 0, corresponding to equal or stronger support for the direction-flipped decoy than for the true mechanism direction.

#### Magnitude failure

For key genes with observed effects, we quantified whether the predicted response magnitude was substantially under- or over-estimated. The per-gene log effect-size agreement was

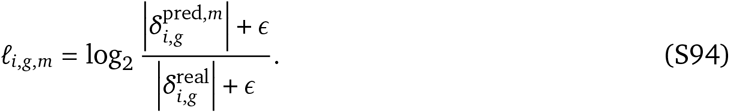

The case-level magnitude error was

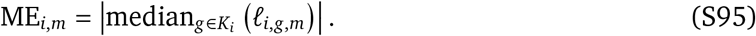

Magnitude failure was defined as ME_*i,m*_ *>* 1, equivalent to a median key-gene effect-size deviation greater than twofold on the absolute Hedges’ *g* scale.

#### Generic-response shortcuts

To determine whether a prediction supported a training-derived generic drug-response program more strongly than the case-specific signature, generic signatures were constructed independently within each data split using only training cases. For each gene, we computed the fraction of training cases in which the true effect exceeded a positive or negative threshold,

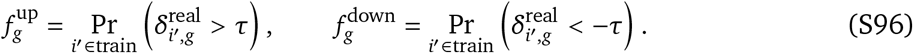

High-frequency upregulated and downregulated genes, or modules obtained by clustering the training effect matrix, were retained as signed generic signatures. For a test case, the true and strongest generic scores were

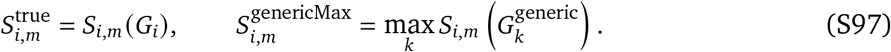

The generic shortcut score was

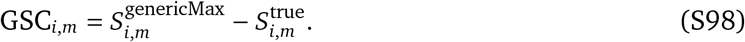

Generic-response shortcut was recorded when GSC_*i,m*_ *>* 0.

#### Non-matching signature decoys

Case-specific signature specificity was tested by comparing the true key-gene signature with decoy signatures curated from other cases. Candidate decoys were required to come from a different case, preferably a different drug, and to have low key-gene overlap with the target case:

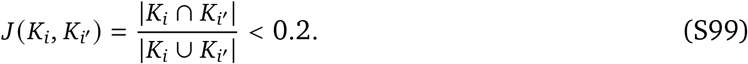

Decoys were further selected to have similar gene-set size and baseline expression distributions in the control cells, with source, cell context, dose, time and study platform matched when possible. For a decoy set D_*i*_, the strongest non-matching decoy score was

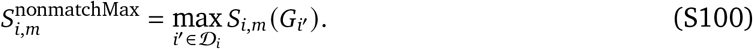

The signature specificity gap was

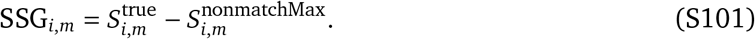

Signature-specificity failure was recorded when 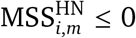 or SSG_*i,m*_ ≤ 0, indicating that the true mechanism genes were not separated from matched random genes or from plausible non-matching mechanism signatures.

#### Strength-matched non-overlapping signature pairs

Finally, we tested whether models distinguished case-specific signatures beyond global perturbation strength. The observed perturbation strength of case *i* was defined as

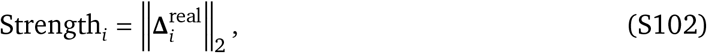

with standardized combinations of alternative strength measures used for sensitivity analyses. For each target case, we selected one or more decoy cases *i′* with similar Strength_*i′*_, a different drug, low key-gene overlap and, where possible, matched source, cell context, dose and time. Both the target and decoy signatures were scored using the prediction for the target case. The strength-controlled signature gap was

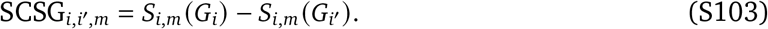

A global-strength shortcut was recorded when 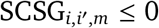. At the model or split level, discrimination was summarized as

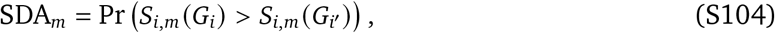

where values close to 0.5 indicate discrimination no better than random among strength-matched pairs.

#### Unified shortcut burden

For each failure mode *k*, we defined a binary indicator *F*_*i,m,k*_ when the corresponding hard-negative criterion was evaluable. The model-level burden for mode *k* was

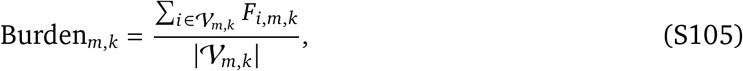

where *V*_*m,k*_ denotes the cases evaluable for model *m* and failure mode *k*. Overall shortcut burden was the mean burden across the seven prespecified failure modes:

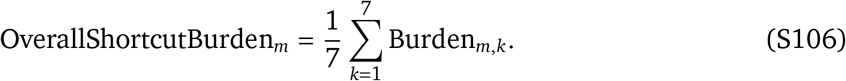

The resulting burden profile summarizes whether poor mechanism fidelity arises primarily from collapse, memorization, direction reversal, magnitude miscalibration, generic-response shortcuts, loss of signature specificity or reliance on global perturbation strength.

